# A scalable method for identifying recombinants from unaligned sequences

**DOI:** 10.1101/2020.11.18.389262

**Authors:** Qian Feng, Kathryn Tiedje, Shazia Ruybal-Pesántez, Gerry Tonkin-Hill, Michael Duffy, Karen Day, Heejung Shim, Yao-ban Chan

**Author notes:** (YBC).

## Abstract

Recombination is a fundamental process in molecular evolution, and the identification of recombinant sequences is of major interest for biologists. However, current methods for detecting recombinants only work for aligned sequences, often require a reference panel, and do not scale well to large datasets. Thus they are not suitable for the analyses of highly diverse genes, such as the *var* genes of the malaria parasite *Plasmodium falciparum,* which are known to diversify primarily through recombination.

We introduce an algorithm to detect recombinant sequences from an unaligned dataset. Our approach can effectively handle thousands of sequences without the need of an alignment or a reference panel, offering a general tool suitable for the analysis of many different types of sequences. We demonstrate the effectiveness of our algorithm through extensive numerical simulations; in particular, it maintains its accuracy in the presence of insertions and deletions.

We apply our algorithm to a dataset of 17,335 DBL*α* types in *var* genes from Ghana, enabling the comparison between recombinant and non-recombinant types for the first time. We observe that sequences belonging to the same ups type or DBL*α* subclass recombine amongst themselves more frequently, and that non-recombinant DBL*α* types are more conserved than recombinant ones.

**Author summary:** Recombination is a fundamental process in molecular evolution where two genes exchange genetic material, diversifying the genes. It is important to properly model this process when reconstructing evolutionary history, and to do so we need to be able to identify recombinant genes. In this manuscript, we develop a method for this which can be applied to scenarios where current methods often fail, such as where genes are very diverse.

We specifically focus on detecting recombinants in the *var* genes of the malaria parasite *Plasmodium falciparum*. These genes influence the length and severity of malaria infection, and therefore their study is critical to the treatment and prevention of malaria. They are also highly diverse, primarily because of recombination. Our analysis of genes from a cross-sectional study in Ghana study show fundamental relations between the patterns and prevalence of recombination in these genes and other important biological categorisations.

## Introduction

Recombination, the exchange of genetic materials between two molecular sequences, is a fundamental evolutionary process in viruses, prokaryotes, eukaryotes, and even between kingdoms [1]. The biological mechanisms of recombination, which differ across different species, lead to the creation of novel ‘mosaic’ sequences in which different regions have distinct evolutionary histories [2–4].

In human population genetics, recombination plays a central role in shaping the patterns of linkage disequilibrium, and thus recombination identification is of importance for estimating recombination rates, quantitative trait loci and association studies [5,6]. Recombination also explains a considerable amount of the genetic diversity of human pathogens [7–9], such as malaria [10] or protozoan parasites [11,12]. It plays a central role for parasites to escape from host immune pressures, or adapt to the effects of antiparasitic drugs. Therefore, the characterisation of recombination events is critical to the clinical treatment and prevention of such diseases.

In phylogenetics, recombination breaks a central assumption, that evolution is tree-like. Not acknowledging recombination can result in severely misleading inferred phylogenies, e.g., the overestimation or underestimation of branch lengths [13–15]. This can be mitigated by the application of phylogenetic network reconstruction methods [16]; however, these methods are still in their infancy. An accurate identification of recombinant sequences would benefit these methods.

Many methods have been developed for identifying recombination events and/or recombinants [3,5,17–20]. They can be roughly characterised into four paradigms:

1. Distance-based methods [1,21–24] look for inversions of distance patterns among the sequences. They usually employ a sliding-window approach to estimate distances and are generally computationally efficient.
2. Phylogenetic methods [8,25–30] look for discordant topologies in adjacent sequence segments, which is taken as a sign of recombination.
3. Compatibility methods [2] test for phylogenetic incongruence on a site-by-site basis. This type of method can be biased by many closely related sequences.
4. Substitution distribution-based methods [31–34] use a test statistic to examine the adjacent sequence segments.

Nearly all available methods require a multiple sequence alignment, which is commonly available for population genetic datasets which have relatively low intra-population diversity. Likewise, many methods (e.g., [21,23,29,30,35]) require a reference panel of known non-recombinant sequences, which potential recombinants can be compared against. In the absence of both an alignment and a reference panel, the available methods for detecting recombinants are limited. Finally, most of the available methods do not scale well to very large datasets.

We focus on the specific application of detecting recombinants in the *var* genes of the malaria parasite *Plasmodium falciparum*. These genes express the *Plasmodium falciparum* erythrocyte membrane protein 1 (PfEMP1), which is the main target of the human immune response to the blood stages of infection. PfEMP1 is expressed on the surface of infected red blood cells and serves to bind host endothelial receptors [36]. It is therefore crucial for the successful proliferation and transmission [37,38] of P *falciparum*. The *var* genes are a large gene family (up to 60 copies per genome) [39], and high levels of diversity in the *var* genes have been observed in a single parasite genome, as well as small local populations [40–44]. This diversity is driven primarily by recombination [10], and so an accurate identification of *var* recombinants is critical to understanding the evolution of the system.

Briefly, the study of *var* genes has revealed a strong domain structure, including multiple Duffy-binding like domains (DBL*α*, *β, δ, ε,* γ, and x) and cysteine-rich interdomain regions (CIDRα, β, γ) [45]. The structure of the gene itself is highly variable in both the number and the composition of these domains. Population genetic studies of *var* genes have focused on sequencing the DBL*α* domain, which almost always appears exactly once in a *var* gene. This domain has been found to be immunogenic [46] and is crucial to understanding acquired immunity and potential for vaccination [47]. Unfortunately, the DBL*α* domain is highly variable, with many thousands of disparate sequences identified. This prevents the construction of a reliable multiple sequence alignment, let alone a phylogenetic tree, and so little is known about their evolutionary history.

Recent evidence [4,47,48] suggests that recombination is uniformly distributed throughout the DBL*α* domain. The first systematic attempt to map out recombination in this domain was performed by Zilversmit *et al.* [4], who developed a method based on a jumping hidden Markov model (JHMM) to align a sequence to its nearest relations in a reference dataset, allowing jumps between sequences which represent recombination events. They used this method to “paint” each sequence according its nearest relations. This was further exploited by Tonkin-Hill *et al*. [48], who studied a large dataset of *var* genes around the world. They found a strong geographic population structure among the genes coming from different countries.

Although these works were valuable in uncovering the recombination structure of *var* genes, there is still much work to be done. The method of Zilversmit *et al*. does not identify recombinant sequences, only recombination events; by identifying the sequences themselves, we can investigate the differences between the recombinants and non-recombinants, and thus determine the effect of recombination on the structure and function of the gene. However, the diversity of the sequences and lack of an alignment and reference panel make it difficult to apply current methods for this task.

In this paper, we develop a new method to identify recombinants in a large dataset of unaligned sequences. This method exploits the information produced by the JHMM method, combining it with a distance-based comparison to identify recombinants. We have applied this method to a large dataset of DBL*α* sequences, producing several new biological results concerning the patterns of recombination in this domain. Extensive simulations also confirm the accuracy and applicability of our method.

## Methods

We propose a novel method to detect recombinant sequences in a set of unaligned protein or DNA sequences. This method is specifically designed to handle sequences for which it is difficult to construct a multiple sequence alignment. It takes as input a set of homologous sequences, and outputs the sequences that are identified as recombinant, their putative parents, and the corresponding breakpoints. Note that extant sequences are identified as the ‘parents’ of the recombinant; more accurately, we identify the descendants of the ancestral sequences which were the parents of the recombination.

Our method combines several previous methods (the JHMM method of [4] and the MAFFT algorithm of [49]) with a novel distance-based approach to identify recombinant sequences. The method consists of the following steps; see Fig 1 for a graphical overview.

1. We apply the JHMM method of Zilversmit *et al.* [4] to represent each sequence as a ‘mosaic’ of segments from other sequences in the dataset.
2. From the mosaic representations, we identify triples of segments which contain a recombinant segment and its two parents. The mosaic representations provide pairwise alignments for each of these triples, which we then complete to three-way alignments with the MAFFT algorithm [49].
3. Using a distance-based approach, we identify the recombinant sequence in each triple.

**Fig 1.**
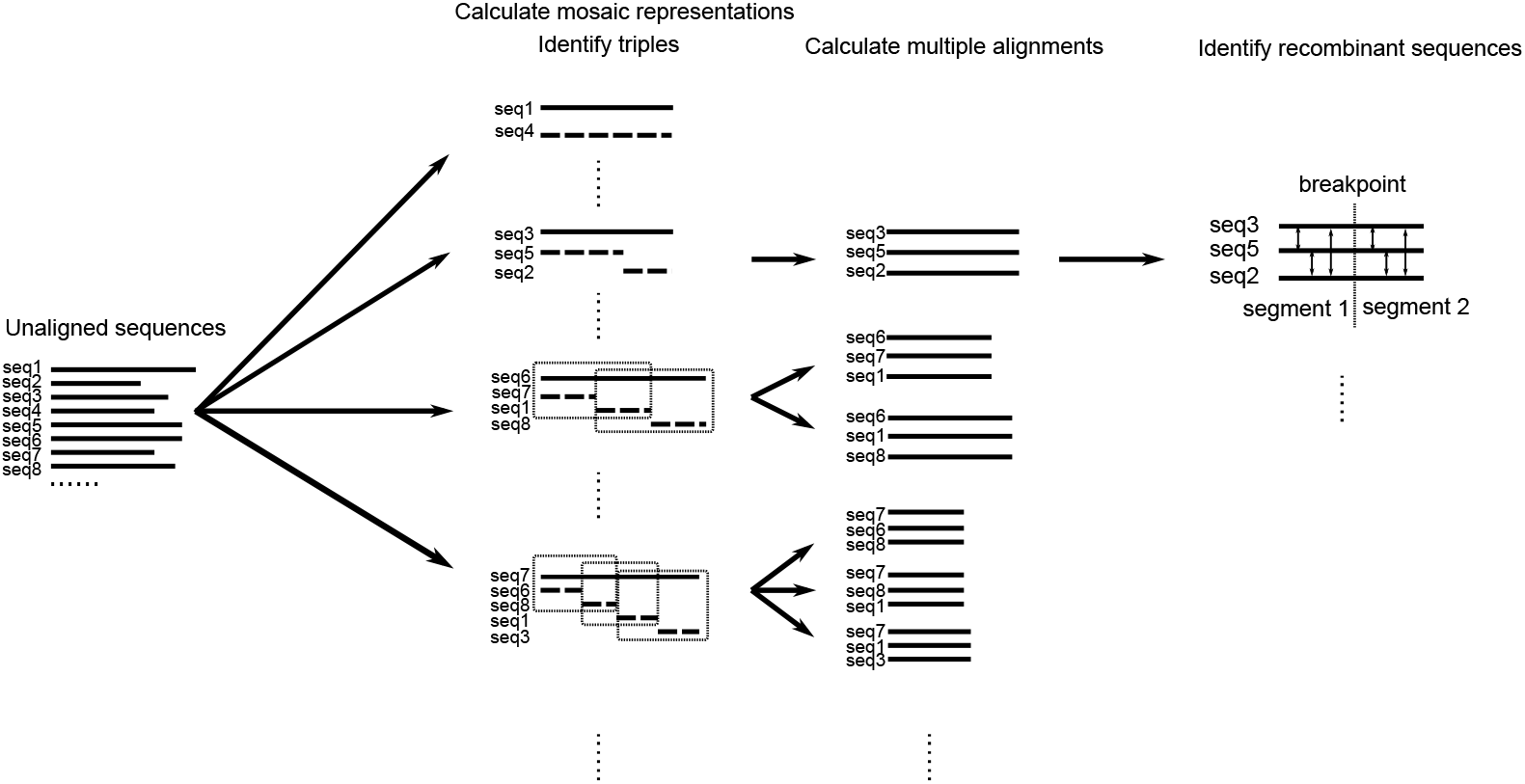
A schematic of the algorithm. From an input set of unaligned sequences, we first use the JHMM method to represent each sequence as a mosaic of other sequences. Next, we identify triples of segments, consisting of a recombinant segment and its two parents, and complete their alignment with the MAFFT algorithm. Finally, we identify the recombinant in each triple using a distance-based approach.

We discuss each step in detail in the following sections.

## Calculating mosaic representations

In this step, we use the jumping hidden Markov model of Zilversmit *et al.* [4] to express each sequence as a ‘mosaic’ combination of the other sequences in the dataset. This model was designed to uncover the patterns of recombination in a set of unaligned sequences.

In this model, each character in a ‘target’ sequence is considered to be a copy from a character in a sequence in a reference set (‘source’ sequences). The hidden state of the Markov model is the (position of the) character which is copied. The copy may be imperfect, representing mutation. After a character is copied, the next character in the target sequence is usually copied from the next character in the same source sequence. However, with small probabilities:

- the source character may switch to any character in any position in another sequence, representing recombination;
- the model switches to an ‘insertion’ state, where the target character is chosen randomly and the source character does not move;
- the model switches to a ‘deletion’ state, where the source character moves forward without being copied.

If the models is in an insertion or deletion state, it continues in this state until (with a small probability per character) we return to copying characters from the current source sequence.

We note that this model is descended from the seminal HMM of Li and Stephens [6], which has seen wide usage in many different applications involving recombining sequences. This model is largely similar, but only works on aligned sequences, and recombination can only switch between characters in the same position in the alignment. This restriction results in a more efficient model with fewer hidden states, but one which cannot be used for unaligned sequences.

We use the Zilversmit *et al.* model here by taking each sequence in our dataset in turn as the target sequence and using every other sequence in our dataset as the source sequences. We first estimate the parameters of the model, following Tonkin-Hill *et al.* [48]. The parameters are the probability of gap initiation *δ*, the probability of gap extension *ε*, and the probability of recombination *ρ*. We first set *ρ* to zero, and compute maximum likelihood estimates for *δ* and *ε* with the Baum-Welch algorithm (see [50]). We then calculate the composite likelihood of all sequences for all values of *ρ* over the interval [0,0.1] under the estimated 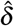 and 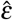, and choose the value of ρ which maximises this likelihood as our estimate 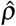.

Finally, we calculate the Viterbi path for each target sequence to find the most probable sequence of hidden states (copied characters, insertions, and deletions). The result is a ‘mosaic’ alignment for each sequence to a series of segments from the other sequences in the dataset. An example of this can be seen in [4, Figure 2A].

**Fig 2.**
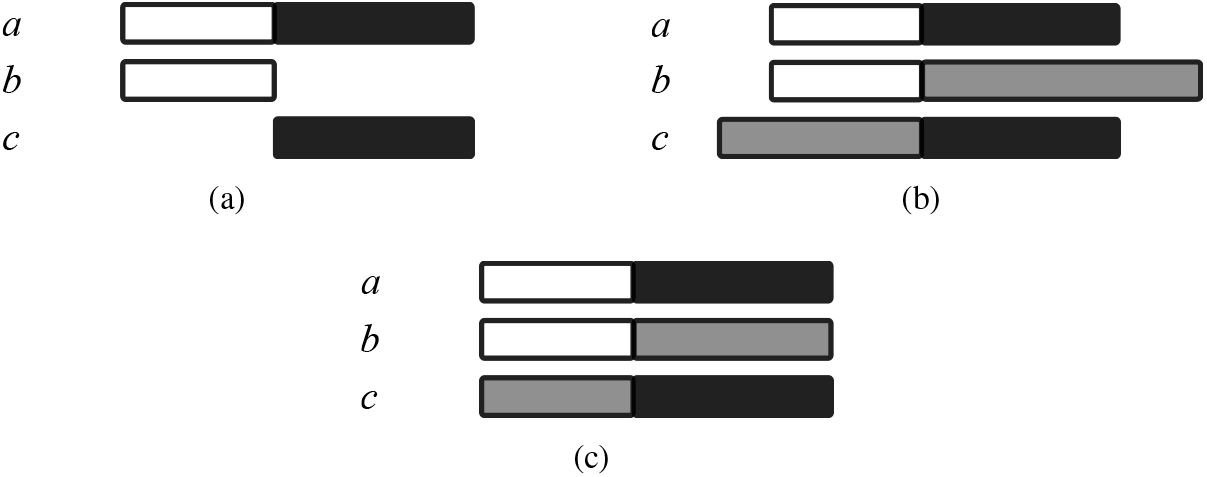
An example of calculating a multiple sequence alignment with MAFFT. (a): A segmental pairwise alignment generated by the JHMM method. Segments from sequence *a* are aligned to segments from sequences *b* and *c* respectively. (b): Using MAFFT, we include the corresponding segment from the third sequence into the pairwise alignment on either side of the breakpoint. (c): By trimming the alignments, we generate a multiple alignment.

For large-scale datasets, training the JHMM model is a significant bottleneck for our method. We again follow [48], and use the Viterbi training algorithm [51] in place of the Baum-Welch to estimate δ and ε, and calculate the composite likelihood over 1000 randomly selected sequences to estimate ρ. This allows us to analyse large datasets (such as the DBL*α* dataset in Section “Analysis of DBL*α* sequences from a cross-sectional study in Ghana”) in a practical timeframe with only a small loss in accuracy.

## Identifying recombinant triples and calculating multiple sequence alignments

For each sequence, the JHMM method produces an alignment of that sequence to segments from the other sequences. Whenever the source segment changes, we consider this to represent a recombination event at that breakpoint. It is not necessarily the case (see below) that the target sequence in this case is the recombinant sequence, and the two source segments come from the parents of the recombination. However, we do know that the target sequence and the two source sequences form a ‘recombinant triple’, that is, are the two parents and the child of a recombination.

Therefore, for each breakpoint in each sequence, we identify the triple of the target sequence and the two sequences which contain the source segments before and after the breakpoint as a recombinant triple. We do this for all target sequences, resulting in a list of recombinant triples, some of which may refer to the same recombination event. Sequences which do not infer a breakpoint do not generate any triples.

We will apply a distance-based method to these triples to identify the true recombinant sequence for each one. To calculate distances, we require a multiple alignment of the segments from these three sequences. However, the JHMM method only provides a pairwise alignment of each target segment to one source segment. We take these pairwise alignments and add the corresponding segment from the remaining source sequence in the triple, using the MAFFT algorithm [49]. For each triple, this results in a multiple alignment of the segments surrounding the breakpoint. See Fig 2 for an overview of this process.

Note that we require a sufficient sequence length on either side of the breakpoint in order to calculate distances accurately. Moreover, we observe in practice that short source segments resulting from the JHMM method tend to be artifacts of the method, rather than representing multiple consecutive recombinations (see S1 Fig). To address this, we exclude triples for which the aligned segment on either side of the breakpoint is less than 10AA, which we found to be a suitable threshold in practice.

## Identifying recombinant sequences

### Identifiability: a phylogenetic perspective

The main novelty in our method is the ability to identify which member of a triple is the true recombinant. It is important to note that the JHMM method does *not* identify the recombinant, but instead finds the (segments of) extant sequences which are the most closely related to the target sequence.

This can be illuminated by considering an explicit phylogenetic network [16] with three aligned sequences and one recombination as an example, as shown in Fig 3. Here, we can translate a phylogenetic network to the corresponding mosaic representations, assuming the JHMM method estimates the distances between sequences perfectly. It can be seen that the same mosaic structure can result from networks with different recombinants.

**Fig 3.**
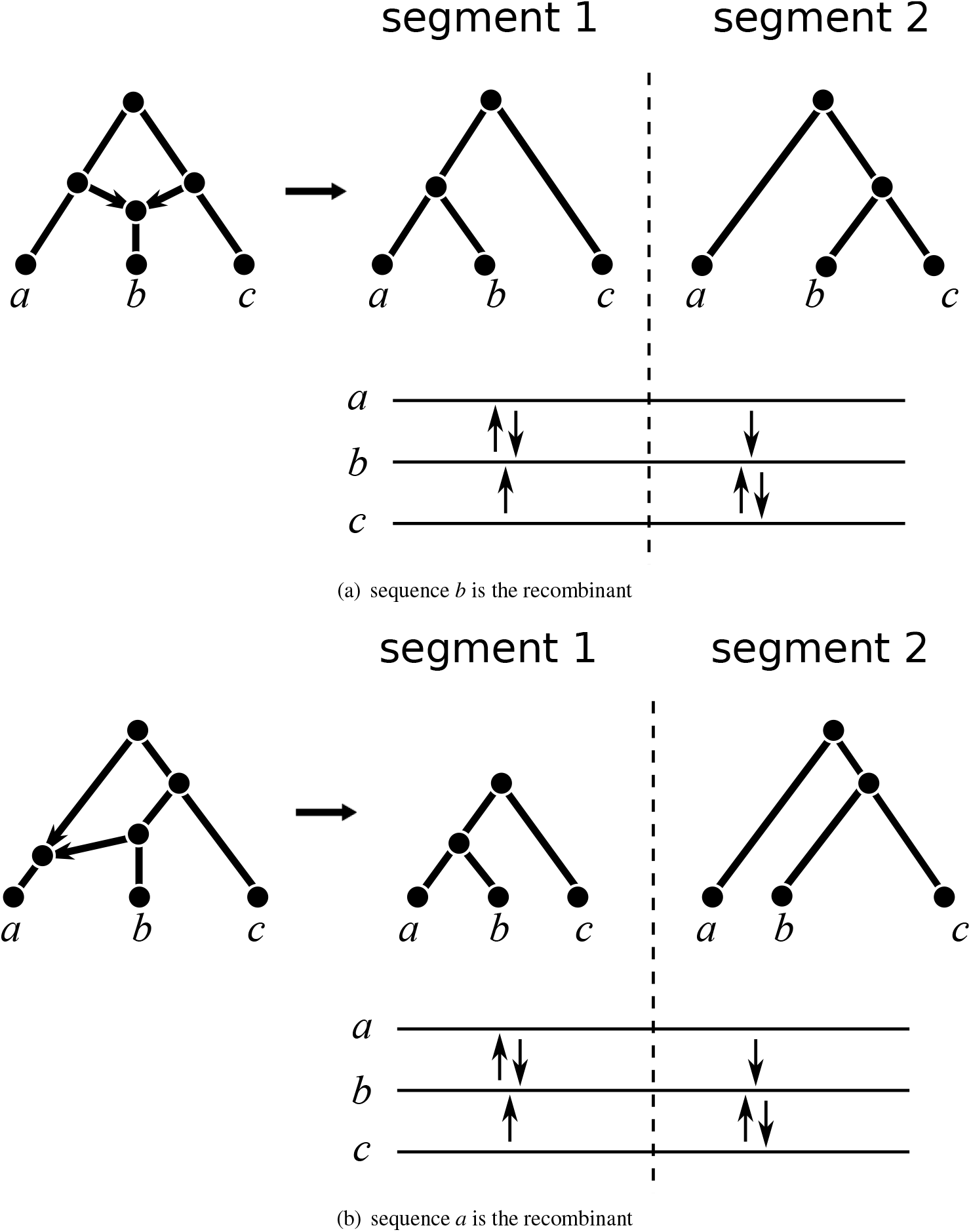
Identifiability of networks from the JHMM output. Here, two networks with different recombinants produce the same profile tree topologies, and thus the same JHMM output. The JHMM output is depicted below the profile trees, with arrows from each target segment pointing to the matching source segment (so, for example, if *b* is the target sequence, it is matched to source sequence *a* in segment 1 and *c* in segment 2 in both cases). Both cases produce identical JHMM output: in particular, sequence *b* is matched to two different source sequences even though it is not necessarily the recombinant.

In fact, as discussed at length in [52], this is an unavoidable problem with the identifiability of phylogenetic networks; networks cannot be distinguished solely by the topologies of displayed trees, which the output of the JHMM method is dependent on. The solution, as given in [52], is to use (inferred) branch lengths to distinguish between the networks, and thereby identify the recombinant.

When the phylogenetic network only consists of three sequences and one recombination (as in Fig 3), it is easy to translate the network to the JHMM output, and thus use it to find the recombinant. However, the problem rapidly becomes much more complicated with more sequences and/or recombinations, and indeed for ancestral recombinations (predating a divergence) it’s not even clear how to define an extant ‘true recombinant’. To avoid this problem, we only identify triples of sequences as in Section “Identifying recombinant triples and calculating multiple sequence alignments”, and assume that only one recombination occurs in the recent evolutionary history of each triple. For large datasets, we are essentially assuming that recombinations are ‘sufficiently far apart’ either in the network or in the genome that they do not interact with each other.

From a phylogenetic perspective, we can see that when this assumption holds, identifying only triples breaks down a complicated network into repeated cases of a three sequence–one recombination network, for which we can identify the recombinant. See Fig 4 for an example of this.

**Fig 4.**
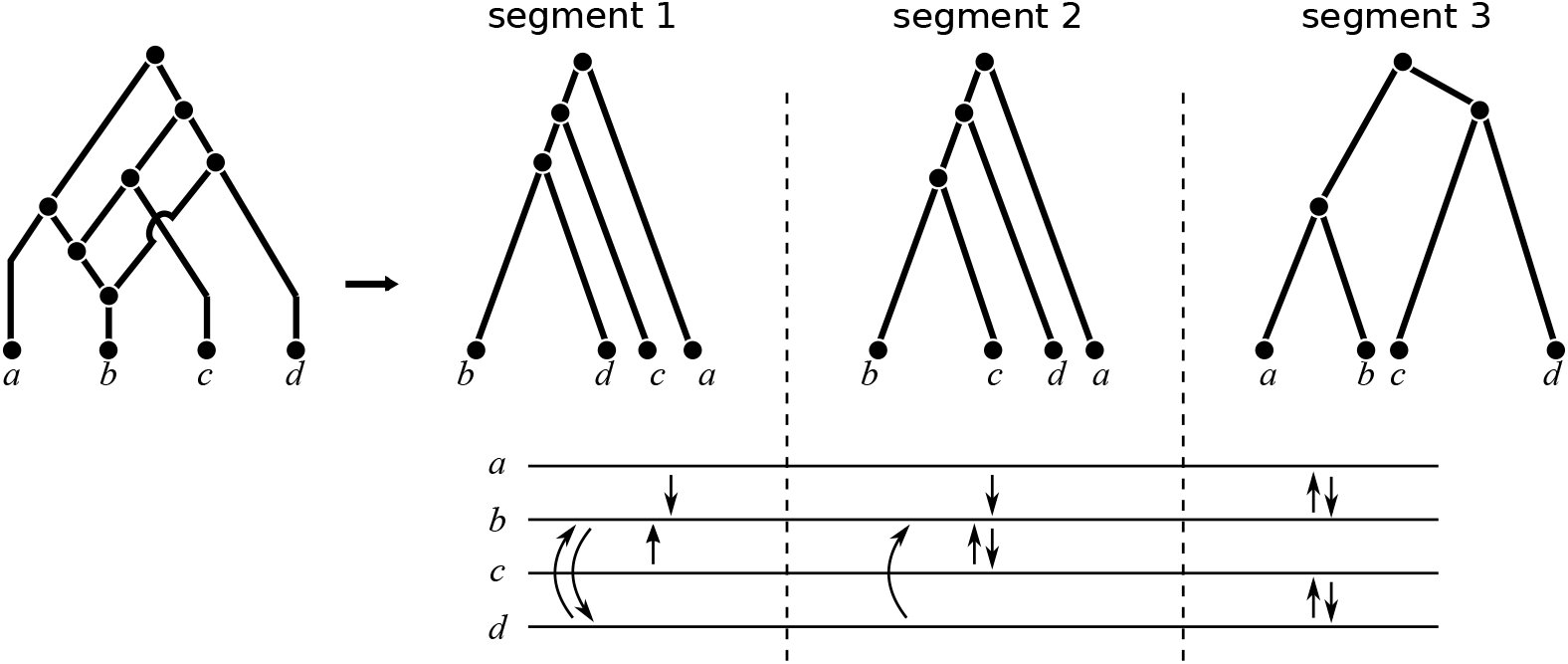
Decomposing a network into triples. At the first breakpoint, the triple {*b, c, d*} is identified from target sequence b, while at the second breakpoint, {*a, b, c*} is identified from sequence *b*, and {*b, c, d*} from sequences *c* and *d*. In all cases, distance-based recombinant identification will obtain the correct recombinant (*b* at both breakpoints).

### Distance-based recombinant identification

Our algorithm is based on the well-known principle [1,17,32,53] that two non-recombinant sequences will have a similar evolutionary distance all along the sequence; that is, the distance between the two sequences does not change before and after a recombination breakpoint in a third sequence. Conversely, the distance between a recombinant sequence and another sequence does change at a breakpoint. Using a distance-based method here allows us to avoid an expensive tree or network inference step and thus scale our method to many sequences.

We thus calculate, for each recombinant triple {*a, b, c*}, the evolutionary distance between each pair of segments before and after the breakpoint. We use here the BLOSUM62 distance [54,55] for amino acids and Hamming (mismatch) distance for DNA sequences (these could in principle be substituted by a large variety of ways to calculate evolutionary distance). We denote these distances by *D*_1_ and *D*_2_ for the first (pre-breakpoint) and second (post-breakpoint) segment respectively.

We then compare the distances for each pair of sequences in the triple before and after the breakpoint; the pair with the smallest absolute difference in distance are inferred to be the two non-recombinant sequences, while the third is inferred to be recombinant. Formally, we have

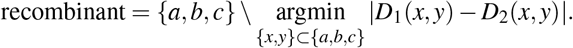

This method identifies one recombinant from each recombinant triple; note that one recombination may generate one or more triples, but the identified recombinant from each of these triples should be the same. We apply this to all triples identified above, generating a list of recombinants in the entire dataset and their putative parents.

## Calculating support values

In addition to identifying recombinant sequences, we can also measure the uncertainty in our identification by using bootstrapping. Bootstrapping in phylogenetics is a standard statistical tool [56], widely used to assign uncertainties to branches on a phylogenetic tree. We use the same basic idea here.

For each multiple alignment of a triple, we resample characters in the alignment (columns) within each segment, with replacement. This provides us with a resampled alignment, and we generate 100 replicates per triple. We then run our distance-based method to identify the recombinant for each replicate. The proportion of replicates which infer the same recombinant as the original alignment is the support value of this detection. The larger the support value, the more certain we are of the detection.

### Efficiency

The complexity of the method is dominated by the first step of estimating the parameters via the Baum-Welch algorithm. As shown in [4], each iteration of the algorithm is *O*(*n*^2^*l*^2^) in time and memory, where *n* is the number of sequences and *l* the length of each sequence. The number of iterations required is not constant, but is generally small (less than 10).

## Results

### Analysis of DBL*α* sequences from a cross-sectional study in Ghana

#### Dataset

We applied our method to detect recombinants and breakpoints in a dataset of DBL*α* sequences collected from individuals with microscopically confirmed *P. falciparum* infections *(isolates)* living in the Bongo District, in the Upper East region of Ghana (GenBank BioProject Number: PRJNA396962) [57,58]. Details on the study population, data collection procedures, and epidemiology have been published elsewhere [59–61]. This dataset consists of 35,591 previously published DBL*α* sequences collected from 161 isolates.

#### Preprocessing

We follow the standard pipeline used in [43,48]. The DNA sequences were first translated into protein sequences, and removed if the resulting sequence contained a stop codon. The protein sequences were then clustered with the Usearch software (v8.1.1861) [62] with a 96% sequence similarity cutoff. The cluster centroids were then taken as a representative sequence for the clusters, which are known as DBL*α types*. This results in a dataset of 17,335 types, each of which may appear in several isolates.

#### Identifying recombinants

We applied our method to this dataset to detect recombinant types. We detected 14,801 (85.4%) of the DBL*α* types to be recombinant.

The analysis was run on a high performance cluster at the University of Melbourne (72 Intel(R) Xeon(R) Gold 6254 CPU cores @ 3.10GHz, 768GB RAM). For estimating parameters, we split the data into 578 subsets of 30 sequences each at every iteration of the Viterbi training algorithm, which were executed in parallel. This was also done for estimating Viterbi paths and identifying recombinants. The time and memory usage is summarised in Table 1. By far the largest bottleneck is the computation of the mosaic representations of the sequences (both parameter estimation and computation of the Viterbi paths); once this was completed, the remaining steps are very efficient even for a dataset of this size.

**Table 1.** Time and memory consumption per subset (30 sequences).

|  | Parameter estimation | Viterbi paths | Recombinant identification |
| --- | --- | --- | --- |
| Time (minutes) | 644.8 | 294.9 | 2.7 |
| Memory (GB) | 21.3 | 21.2 | 0.1 |

### DBL*α* sequences from the same ups type recombine more frequently

The upstream promoter sequences of each *var* gene can be classified into three main ups types, upsA, upsB, and upsC [41]. These ups types (not to be confused with DBL*α* types; instead, they are analogous to DBL*α* subclasses) are associated with disease severity and clinical significance [63], and thus it is crucial to investigate the behaviour of recombinants and recombinations within and between ups types.

Earlier studies on a much smaller dataset [64], based on sequence similarity, proposed that *var* gene recombination preferentially occurs within the same ups type. Using our method, which to our knowledge is the first systematic attempt to detect recombinants in *var* genes in natural parasite populations, we found considerable evidence supporting this hypothesis. Our results are summarised in Table 2.

**Table 2.** Proportions of recombinations from the same ups types. Theoretically expected proportions, based on the base frequencies of the ups types, are given in brackets. All *p*-values are highly significant (< 2.2 × 10^-16^).

|  | Parent-child | Parents | Family |
| --- | --- | --- | --- |
| UpsA vs. upsB/C | 99.7% (85.0%) | 98.9% (85.0%) | 98.5% (77.6%) |
| UpsA, B and C | 85.3% (50.9%) | 65.5% (50.9%) | 51.1% (30.5%) |

Following the method of [41], we classified each DBL*α* type into one of 32 subclasses. The subclasses were then classified into either upsA or upsB/C types (the latter two being difficult to distinguish based on subclasses alone). For greater precision, we also developed a method to distinguish between all three types: we used BLASTP [65] to match each sequence to the closest reference sequence in [41], and then classified that sequence to the ups type of the closest reference sequence.

Having identified recombinant sequences and their putative parents, we then calculated the proportion of recombination triplets which have one parent and the child, both parents, and both parents and the child belonging to the same ups type (‘Parent-child’, ‘Parents’, and ‘Family’ in Table 2). In all cases, we found that the parents and/or the child of a recombination were significantly more likely (p < 2.2 × 10^-16^ from *χ*^2^ tests) to belong to the same ups type. This effect was most strongly noticeable when we divided the sequences only into ups A and B/C types; for example, the two parents and the child were in the same type 98.5% of the time, compared to a theoretical expectation of 77.6%. Similar conclusions were reached when we divided the sequences into three types. Our results strongly reinforce the conclusions of earlier studies, and provide more precision with the division into three ups types.

We also considered the proportions of identified recombinants in each ups type. We found that there was a significant difference in the proportions of recombinants in the three types (*p* = 2.193 × 10^-7^ from a *χ*^2^ test), with upsA having the least proportion of recombinants, and upsC the most (82.3%, 84.9%, and 87.6% from A, B, and C respectively).

### Proportions of recombination differ among DBL*α* subclasses

DBL*α* sequences can be classified according to sequence similarity into 33 subclasses (DBL*α*0.1–24, DBL*α* 1.1–8, DBL*α*2). These subclasses are strongly associated with ups types; however, they also provide greater resolution in dividing the sequences. We thus repeated our earlier analyses with regards to the subclasses.

As with ups type, we found a significant (all *p* < 2.2 × 10^-16^) increase in recombinations with one parent and the child (58.8% vs. 7.9% expected), parents (31.0% vs. 7.9% expected), and both parents and the child (20.6% vs. 1.0% expected) from the same subclass.

We next considered the proportions of identified recombinants in each subclass (Fig 5). We identified seven subclasses (DBL*α*0.1, 5 and 11 were too high, while DBL*α*0.3, 8, 9 and 23 were too low) which were significantly different from the average under a Bonferroni correction for multiple testing. Of particular note is the DBL*α*0.1 subclass, which has been noted to involve more recombinations than other subclasses [10]. We suggest that these subclasses should be explored further to determine if there are some biological factors that may explain these results.

**Fig 5.**
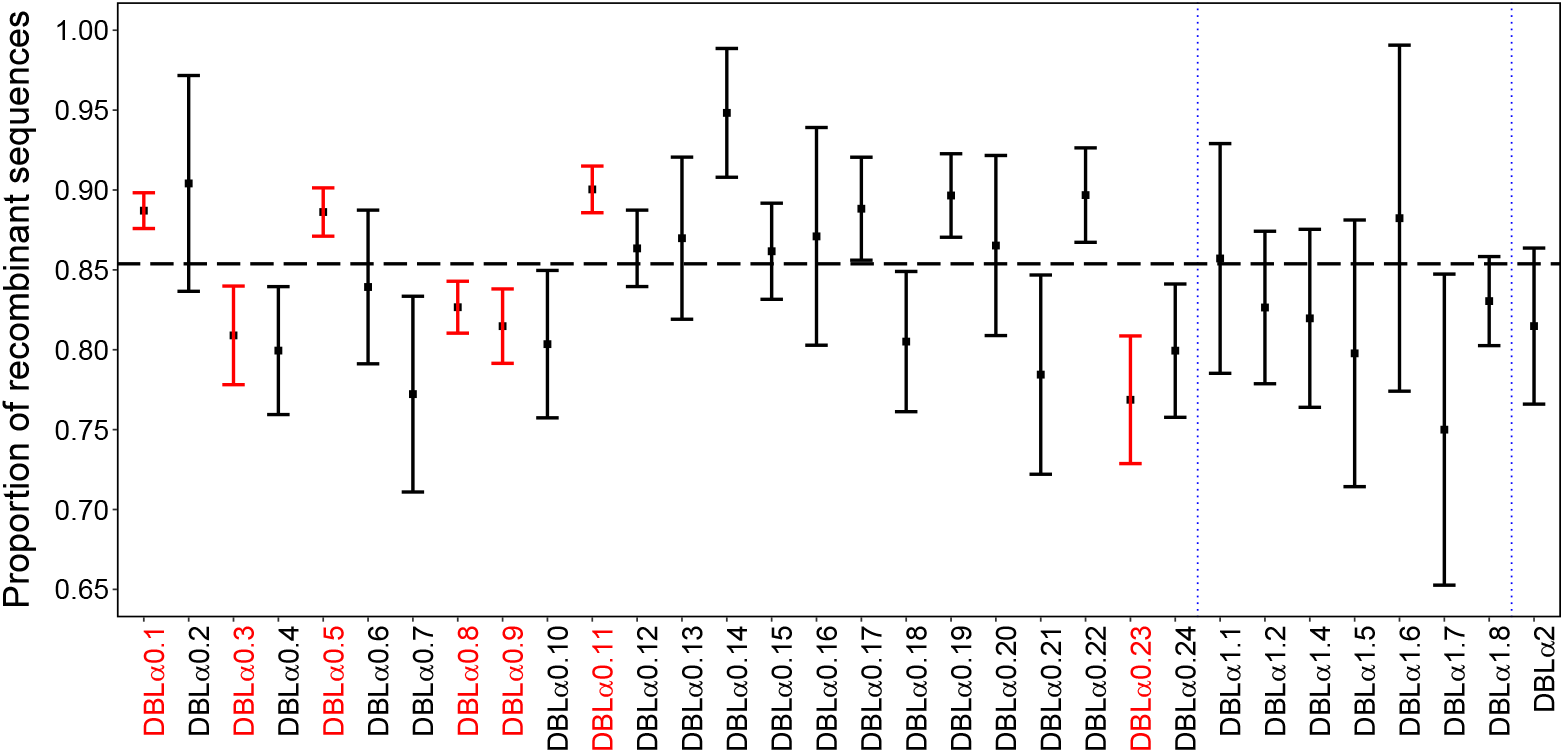
Proportions (and 95% confidence intervals) of recombinants for each DBLα subclass. Subclasses which are significantly different from the overall average are highlighted in red. The horizontal dashed line displays the overall proportion of recombinant sequences in the entire dataset.

We also investigated the proportion of recombinants among individual isolates, and among the two broad catchment areas in the Bongo District (Soe and Vea/Gowrie) that the isolates were collected from. We did not detect any significant differences here, see S1 File and S2 Fig for more details.

### Non-recombinant DBL*α* types are more conserved than recombinant types

It is well known [43,66] that some DBL*α* types are highly conserved, i.e., that they appear in many different isolates. On the other hand, many other types only appear rarely (or even once, in our large dataset). We hypothesise that non-recombinant types are more “stable” than recombinants, and thus may be more highly conserved.

We investigated this hypothesis via the recombinants identified by our method. Firstly, we compared the observed frequencies of the recombinants to the non-recombinants; we found that non-recombinants occurred significantly more often in the dataset (average 4.2 vs. 3.7, *p* = 0.021 from a Wilcoxon rank sum test).

We also considered if there is a difference in the proportions of frequent DBL*α* types in recombinants and non-recombinants. As the frequencies of types are highly right-skewed (see S3 Fig), thus, we thresholded the frequencies at various levels to determine if there were particular frequencies where an effect could be noticed. The results are in Table 3. We found that for a threshold frequency of 5, there were significantly fewer frequent recombinants than non-recombinants; however, this effect becomes less noticeable for larger thresholds. This suggests that there is a high proportion of recombinants which appear very few times in the dataset; these are potentially relatively recent recombinants, which may have not been fixed in the population.

**Table 3.** Proportions of frequent (larger than the threshold) recombinant and non-recombinant DBL*α* types for different thresholds.

| Threshold | 5 | 10 | 15 | 20 |
| --- | --- | --- | --- | --- |
| Recombinants | 17.5% | 4.5% | 2.1% | 1.3% |
| Non-recombinants | 21.0% | 6.0% | 2.3% | 1.6% |
| $P$-value ($\chi^2$ test) | 0.006 | 0.047 | 0.666 | 0.634 |

### Breakpoint positions are associated with homology blocks

It is known that a number of semi-conserved homology blocks (HBs) occur frequently in *var* genes [41]. These HBs recombine at exceedingly high rates [67,68], and are known to be useful in predicting disease severity [36]. We thus investigated the patterns of recombination in DBL*α* types in relation to these homology blocks.

The positions of recombination breakpoints, as found by the JHMM method, are shown in Fig 6. Of particular note is:

- The recombination rate is not constant throughout the sequence, but displays three distinct peaks spaced in roughly equal intervals. These peaks clearly correspond to the three most frequent homology blocks, HB5, 14, and 36, with the height of the peak also corresponding to the frequency of the HB.
- The frequency of breakpoints drops sharply towards either end of the sequence. This is an artifact of the method and does not imply that the recombination rate is lower there; we cannot recognise a recombination which is close to one end of the sequence.

**Fig 6.**
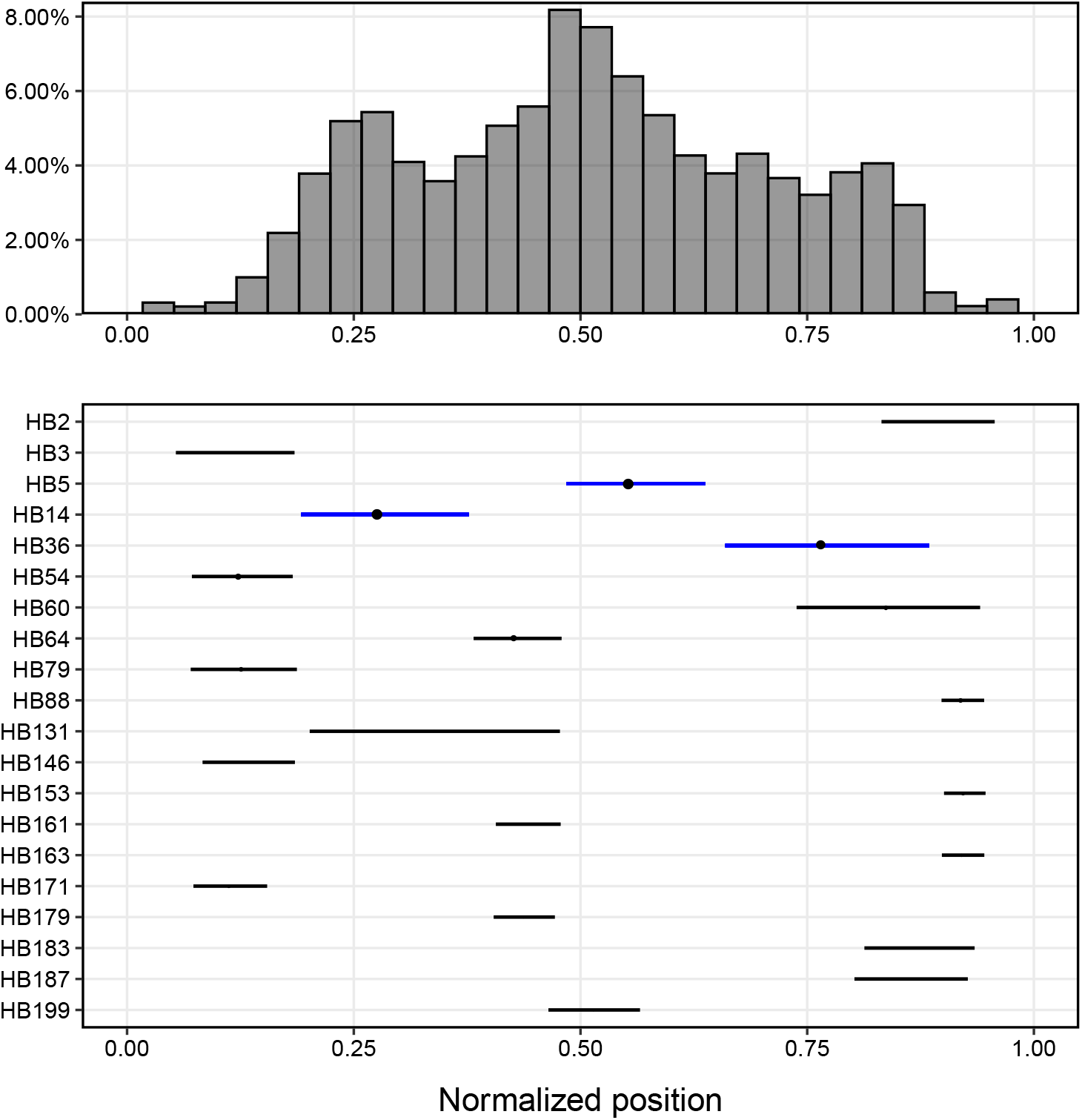
Positions of recombination breakpoints. (Top) The histogram of relative breakpoint positions of recombinations. (Bottom) The position of the most common homology blocks, with circle size proportional to frequency. The three most frequent homology blocks (HB5, 14, and 36) are highlighted in blue.

This reinforces the biological theory that recombination occurs within short identical segments [69].

We also investigated the occurrence of HBs in the recombinant and non-recombinant sequences identified by our algorithm. We discovered that the number of HBs in recombinant sequences were significantly higher than in non-recombinant sequences (5.5 vs. 5.3, *p* < 2.2 × 10^-16^ from Wilcoxon rank sum test). Furthermore, the proportion of sequences containing “important” HBs (5, 14, and 36) were also significantly different between the two groups (83.9% vs. 78.5%, *p =* 1.859 × 10^-11^ from χ^2^ test), indicating that recombinants tend to have more conserved building blocks. Finally, we found that recombinant sequences had higher pairwise HB similarities [36] with each other than non-recombinants (0.629 vs. 0.618, *p* < 2.2 × 10^-16^ from Wilcoxon rank sum test). For more details, see S2 File.

### Simulations

#### Simulation design

We conducted extensive simulations to evaluate the effectiveness of our method. Our simulation protocol is as follows:

1. Simulate a tree (genealogy) under the coalescent (without recombination) using `msprime` [70].
2. Evolve amino acid sequences from a common ancestor along the tree using `Pyvolve` [71]. If insertions and/or deletions are required, we use `INDELible` [72] instead.
3. Generate recombinant sequences from two or more randomly chosen sequences in the dataset, with breakpoints chosen uniformly at random along the genome. The parent sequences are removed from the dataset.

Note that we do not evolve our sequences further after the recombination step; however, since we remove the parents from the dataset, this is indistinguishable from having earlier recombinations in sequences that do not diverge.

In our simulations, we simulate both equal-length sequences (no indels, see Table 4), and unequal-length sequences with indel events (see Table 5), generating unaligned input.

**Table 4.**
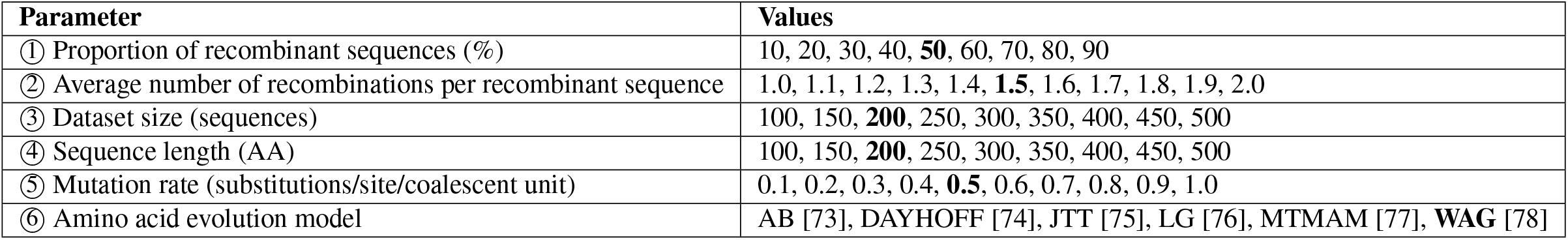
General simulation parameters (no indels). We vary each parameter in turn while holding the others fixed at the default values (in bold).

**Table 5.** Indel simulation parameters (default values in bold). Insertions and deletions are simulated at the same rate, with lengths according to a negative binomial distribution with variance 10.

| Parameter | Values |
| --- | --- |
| ⑦ Indel rate (expected number of indels/site/coalescent unit) | 0.1, 0.2, **0.3**, 0.4, 0.5 |
| ⑧ Mean indel size (AA) | 3.7, 5.2, **6.0**, 6.6, 7.0 |

There are a wide variety of parameters which could potentially affect the performance of the method. Some of these are laid out in Tables 4 and 5. To keep our simulations tractable, we only vary one parameter at a time, keeping the remainder fixed at default values. For each parameter combination, we simulate 100 datasets and run our method on each dataset in turn.

To assess the performance of our algorithm, we calculate the *sensitivity* and *specificity* of our method for each dataset. The sensitivity is defined as the proportion of true recombinants that are correctly detected, while the specificity is the proportion of true non-recombinants that are correctly detected.

### Results

Our results are shown in Figs 7–14. Overall, it can be seen that the method enjoys good performance, with most parameter settings offering both sensitivity and specificity above 70% (and often much higher). We briefly consider the effect of each parameter in turn.

**Fig 7.**
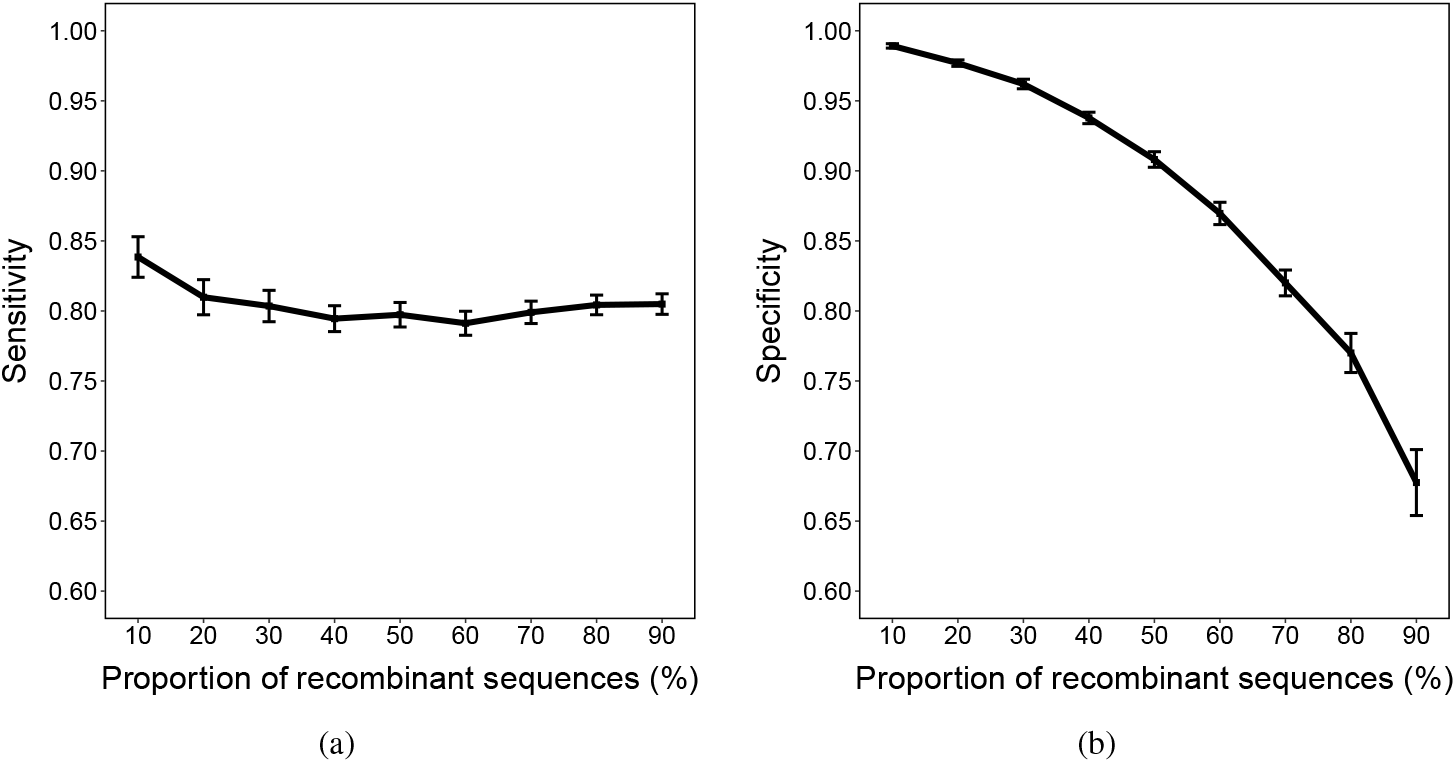
Mean sensitivity and specificity (with 95% confidence intervals) for varying proportions of recombinant sequences.

#### Recombinant proportion

As the proportion of recombinants increases, sensitivity is stable at around 80%, while specificity decreases (Fig 7). Here, more recombinant sequences result (correctly) in a higher number of recombinations detected. It appears that the proportion of true recombinants extracted from the recombinant triples remains largely the same (constant sensitivity); however, there are proportionally more false detections as the number of non-recombinants decreases, resulting in a lower specificity.

#### Number of recombinations per recombinant

As shown in Fig 8, the datasets where there are more recombinations per recombinant sequence appear to have a higher sensitivity, and slightly lower specificity. As for recombinant proportion, an increase in the number of recombinations results (correctly) in more inferred recombinations; unlike that case, the number of true recombinants remains the same here. It appears that the ‘extra’ detections are mostly correct, which results in a greater proportion of true positives (sensitivity increases) and a relatively stable specificity.

**Fig 8.**
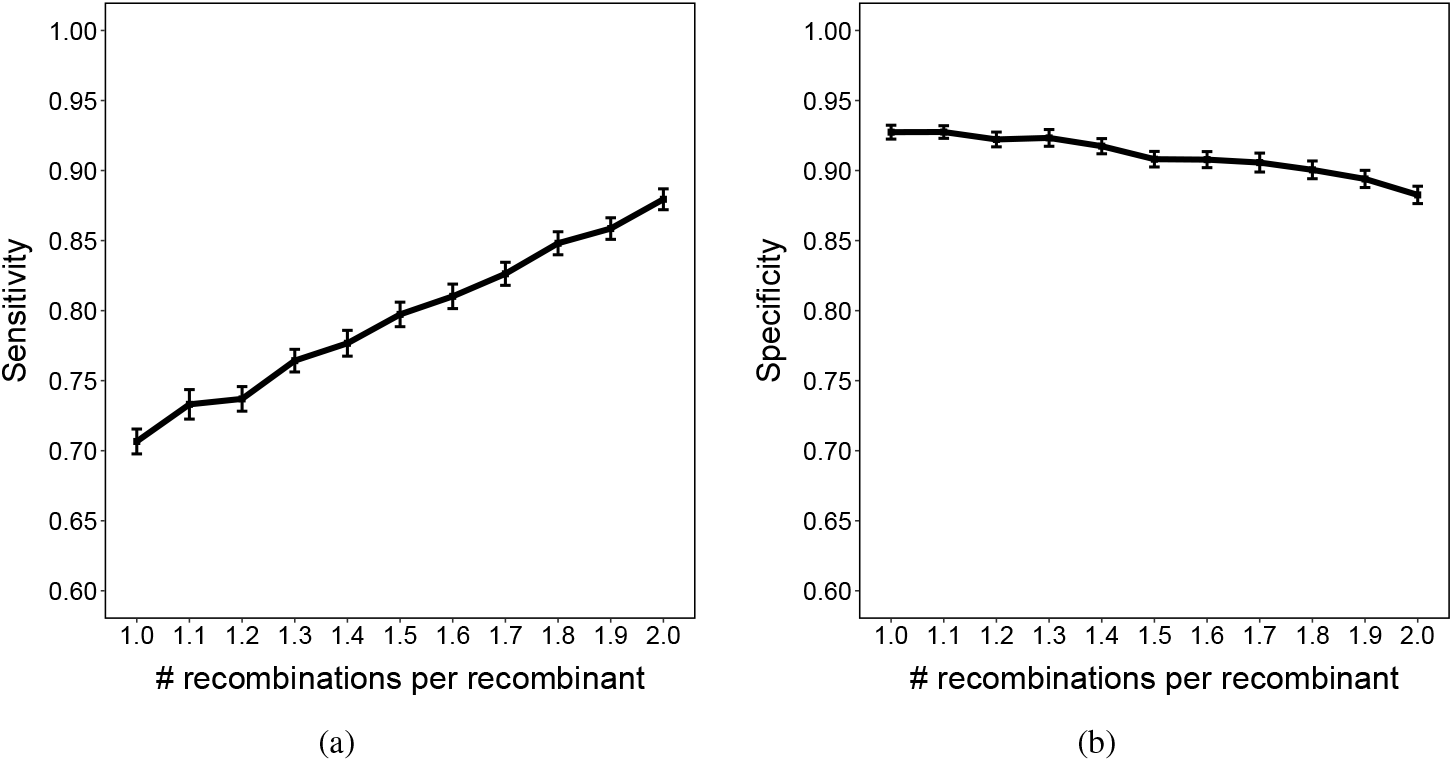
Mean sensitivity and specificity (with 95% confidence intervals) for varying numbers of recombinations per recombinant sequence.

We also conducted a further analysis by matching the distribution of the number of recombinations per recombinant to the Ghana dataset from Section “Analysis of DBL*α* sequences from a cross-sectional study in Ghana” (see S3 File and S4 Fig for more details). Our results indicate that, despite a low specificity (40.0%), a high sensitivity (89.0%) still demonstrates the applicability of our algorithm to real data.

##### Dataset size

Dataset size does not appear to have a drastic effect on the sensitivity of the method, while specificity increases slightly (see Fig 9). It is to be expected that performance increases slightly as information accumulates across a larger dataset, but it is unclear why this is only expressed in the specificity here.

**Fig 9.**
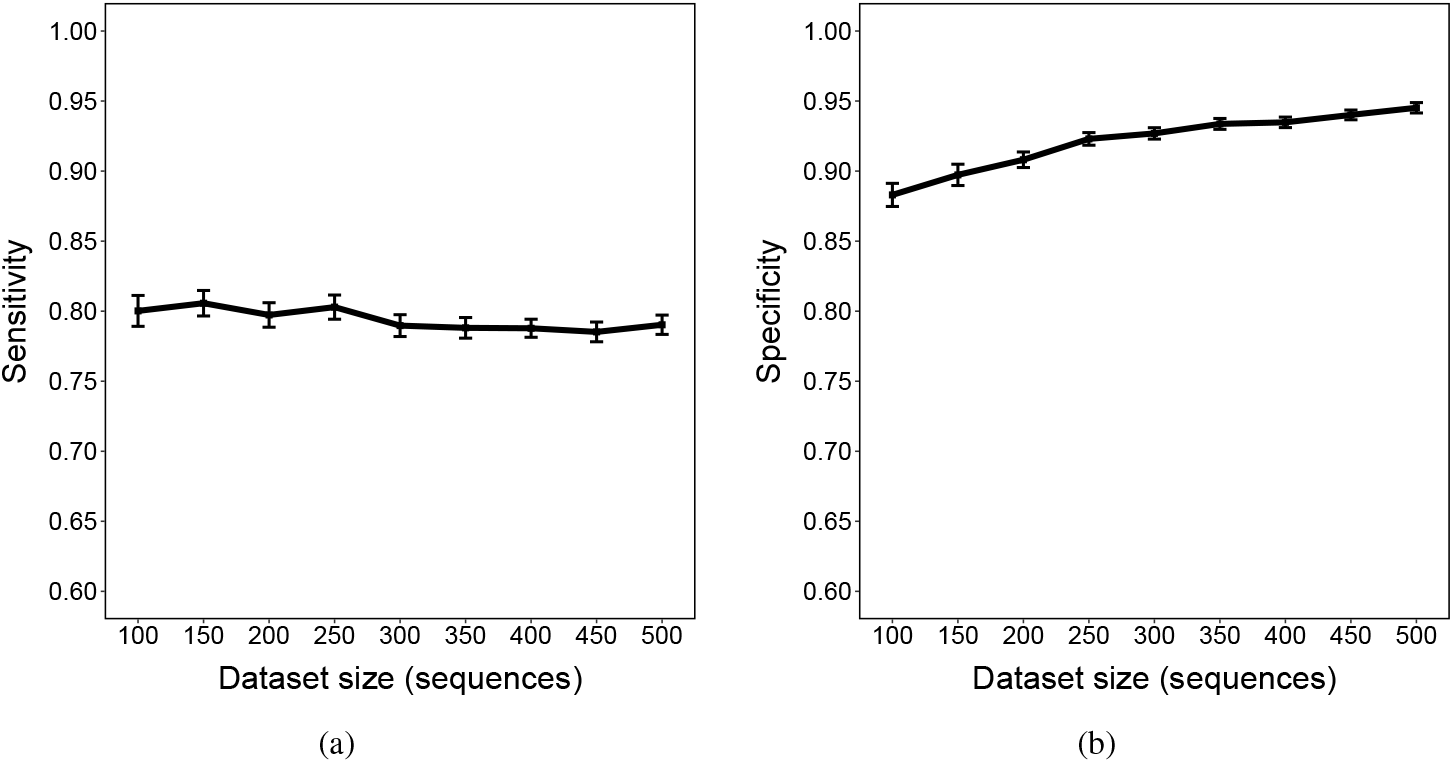
Mean sensitivity and specificity (with 95% confidence intervals) for varying dataset size.

##### Sequence length

Datasets with longer sequence length have much higher sensitivity, and slightly lower specificity (Fig 10). It seems (S5 Fig) that as sequence length increases, the number of recombinations detected also increases, even though the true number of recombinations remains the same. This increase in detections, combined with a fixed percentage of recombinants, results in a effect similar to that seen for the “number of recombinations per recombinant”: an increase in sensitivity and a slightly decreasing specificity.

**Fig 10.**
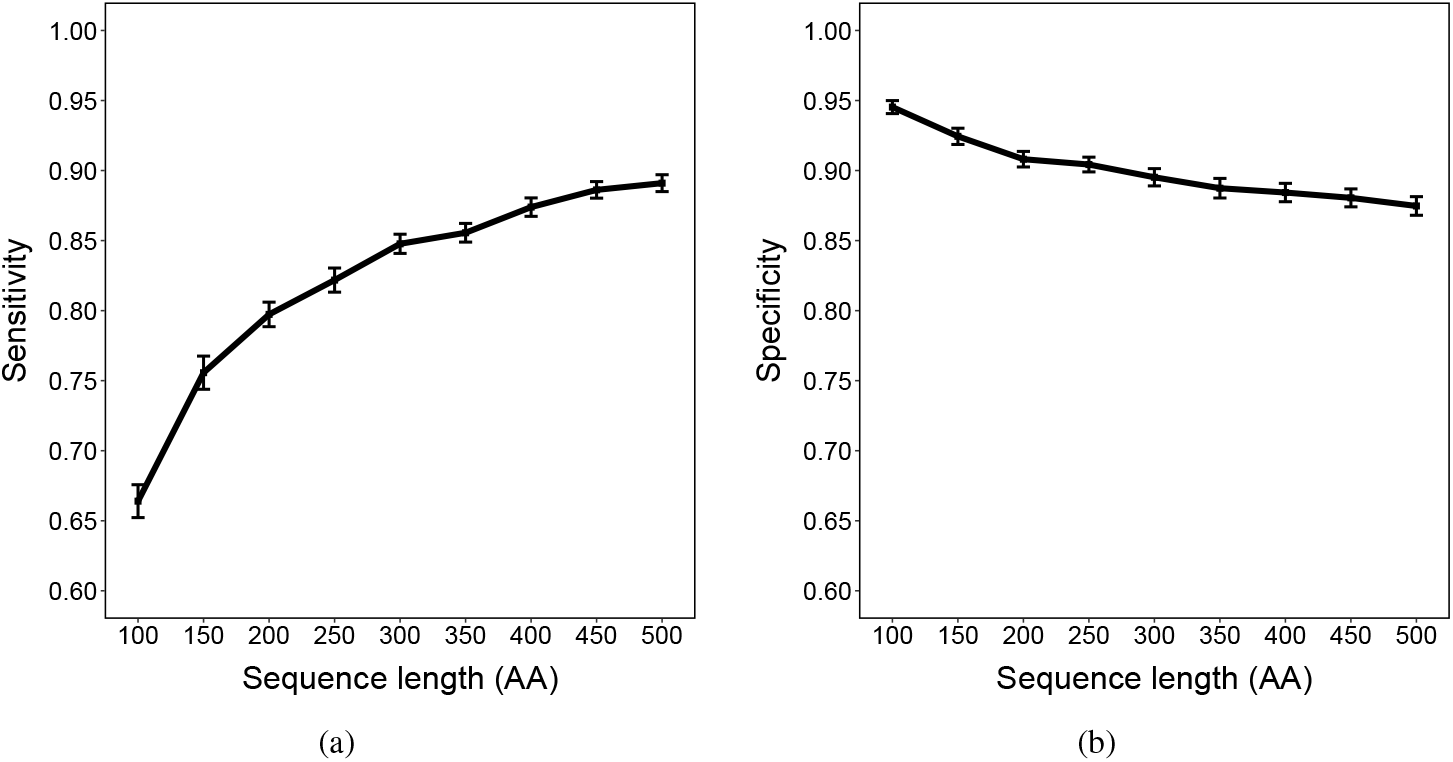
Mean sensitivity and specificity (with 95% confidence intervals) for varying sequence length.

##### Mutation rate

As the mutation rate increases, the sensitivity of the method rapidly increases before levelling out (Fig 11). This makes sense, as if the number of substitutions is too low, the sequences are difficult to distinguish from each other, which makes the results from the JHMM unreliable. Conversely, as the number of substitutions grows, it also becomes more difficult to identify sequences which are closely related to each other, resulting in a decrease in specificity.

**Fig 11.**
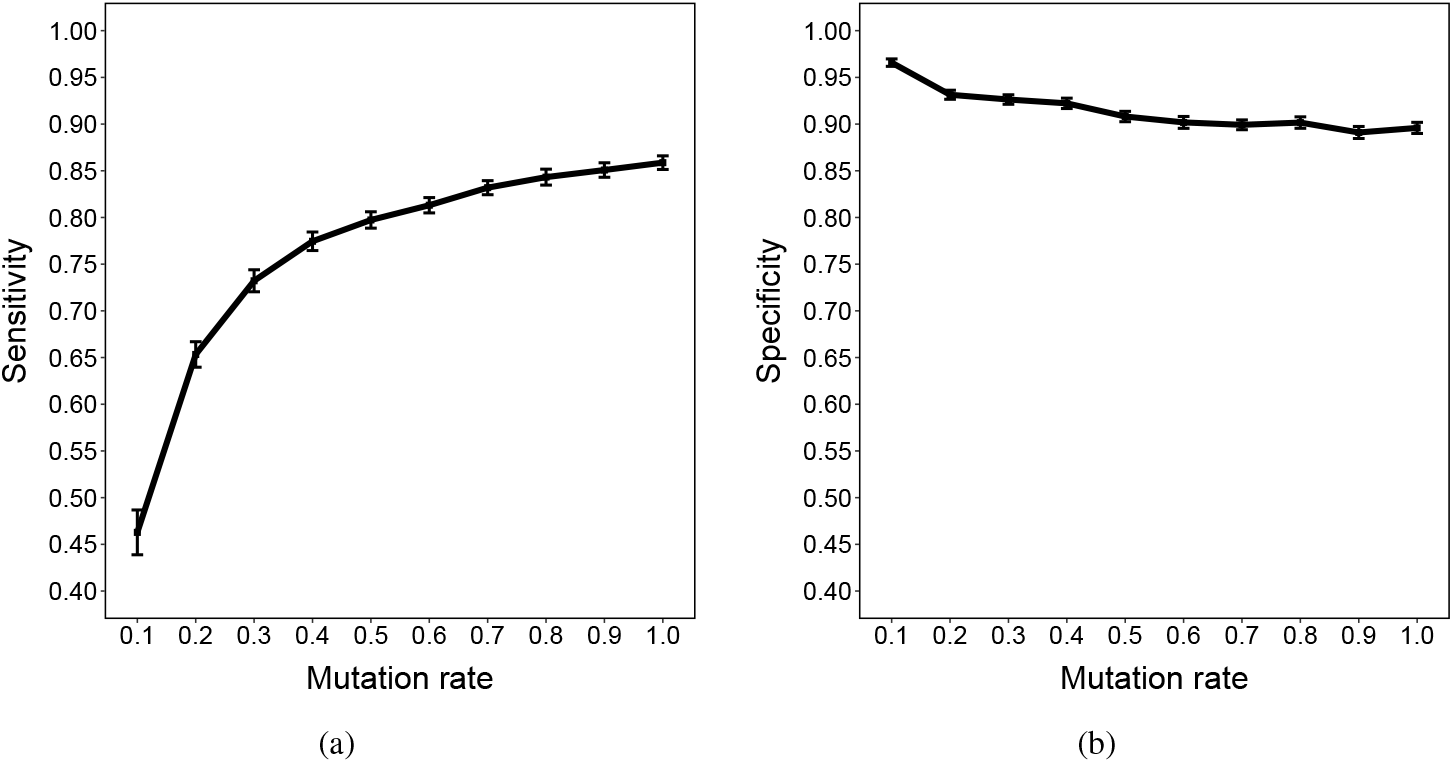
Mean sensitivity and specificity (with 95% confidence intervals) for varying mutation rate.

##### Insertion/deletion parameters

An important feature of our method is its ability to accept unaligned sequences as input. When we include indels in the generating process, we can see (Figs 12, 13) that both sensitivity and specificity remain relatively unaffected, with a moderate decline in specificity as indel rate increases. This indicates that our method is robust to indels even when the indel rate or fragment size is large. In these scenarios, existing methods which only accept aligned sequences would be unable to cope.

**Fig 12.**
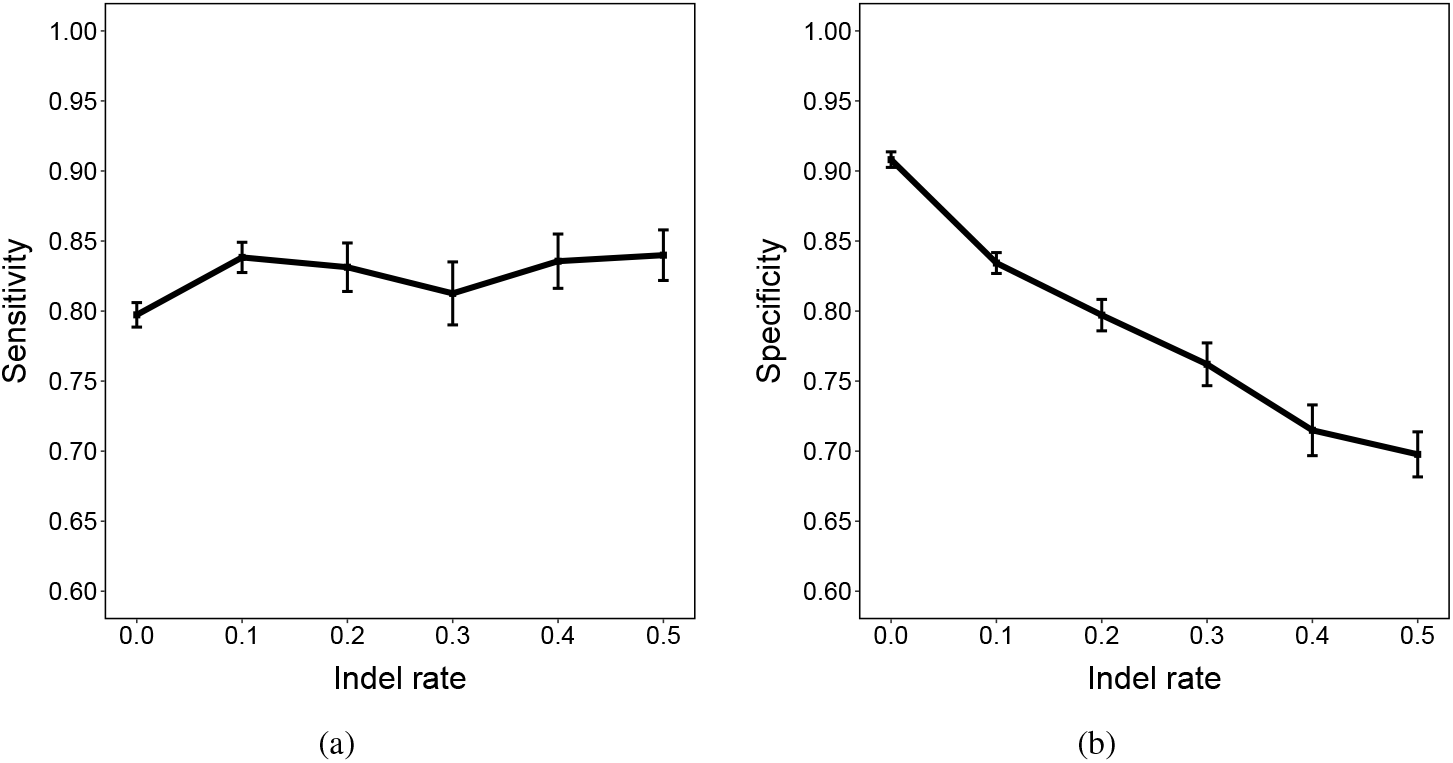
Mean sensitivity and specificity (with 95% confidence intervals) for varying indel rate.

**Fig 13.**
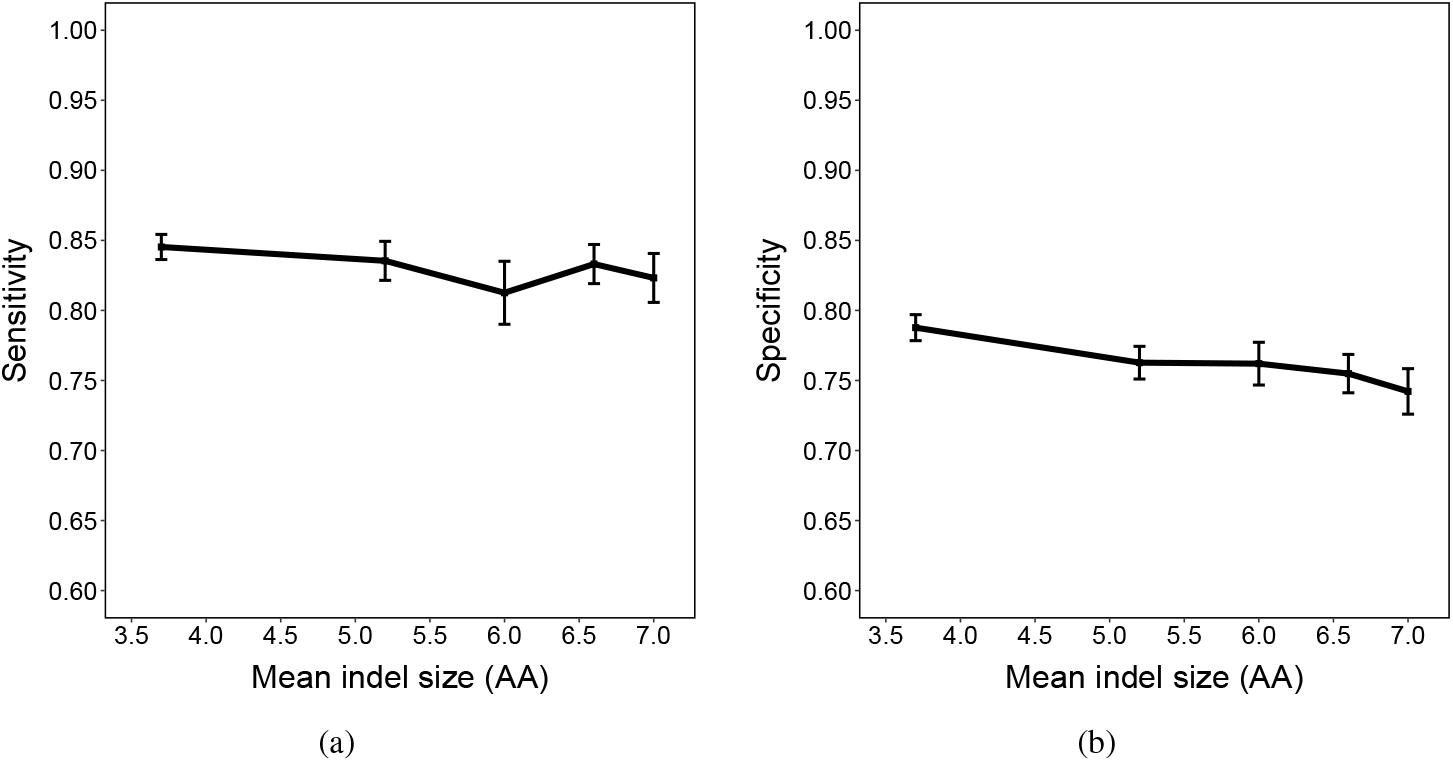
Mean sensitivity and specificity (with 95% confidence intervals) for varying indel size.

##### Other parameters

The method appears to be robust to the stochastic model of amino acid evolution (Fig 14).

**Fig 14.**
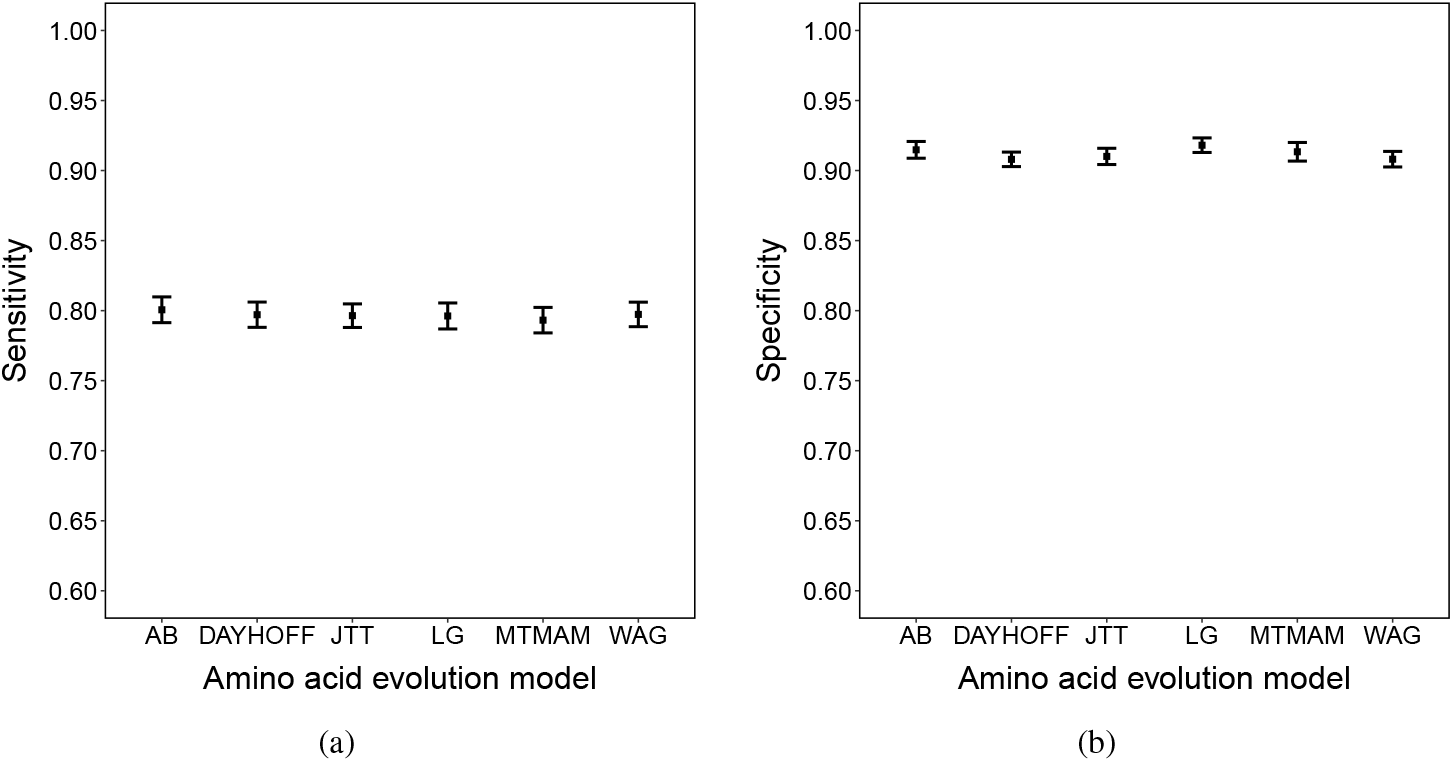
Mean sensitivity/specificity (with 95% confidence intervals) for each model of amino acid evolution.

#### Support values

In addition to detecting recombinants, we also show above how to calculate support values for each detection using bootstrapping. Here, we verify that the calculated values are indeed effective for this purpose. For our simulations, we calculate the support values for each of the correct detections, as well as each of the false positives. The distributions of the support values for the default parameters are shown in Fig 15. Here, we can see that there is a clear separation between the distributions of support values for the true and false positives; while the values for both are relatively high, the support values for true detections are overall much higher. Similar patterns are seen among all the remaining parameter settings (S6 Fig–S13 Fig).

**Fig 15.**
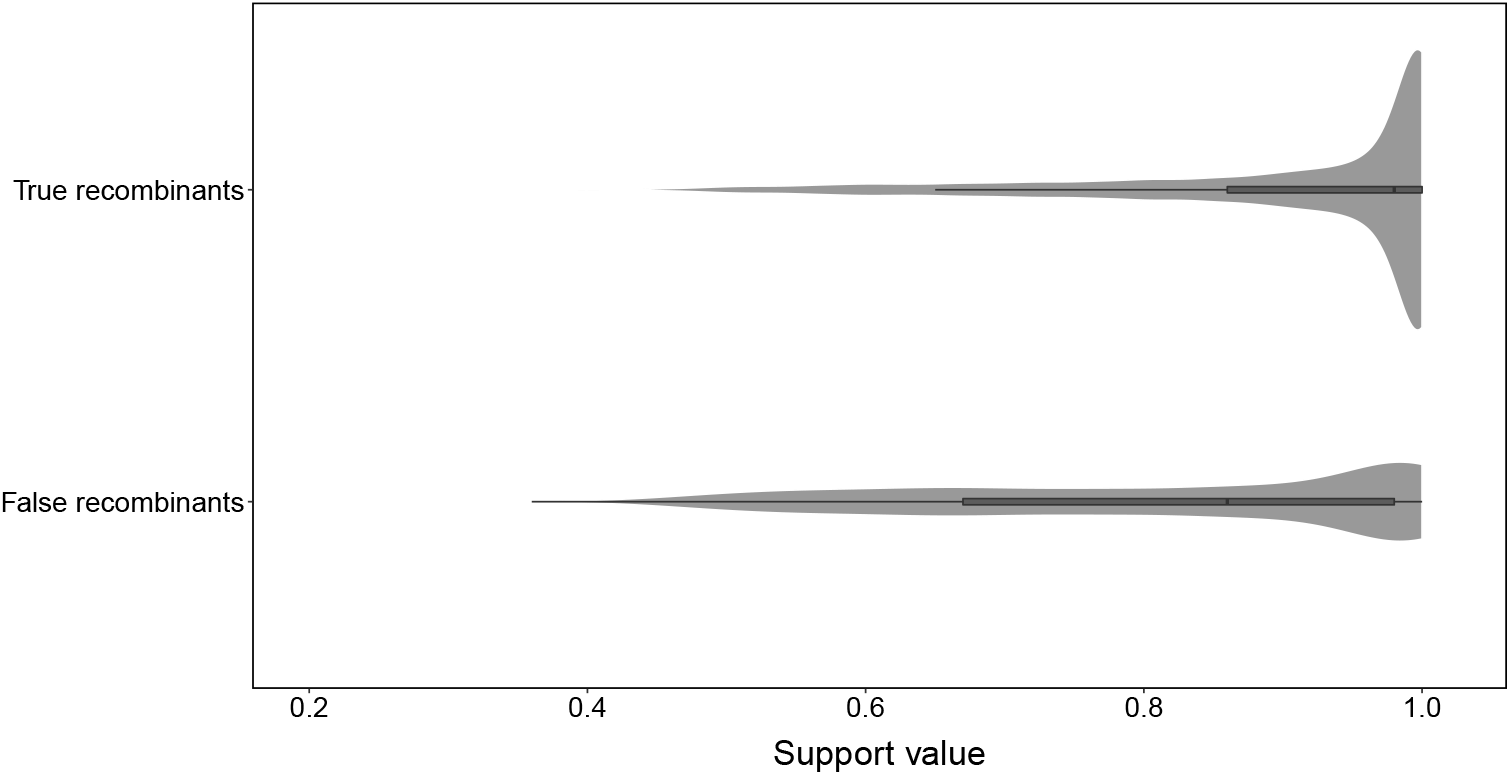
Distributions of support values under default parameters without indel events.

This suggests that we can use a threshold on the support value to refine our detections. This is reasonable if we wish to reduce false positives; however, in practice we found that applying a threshold also reduced true positives (as expected) to an extent which lowered the overall accuracy of the method, so we have elected not to apply it here. Instead, we suggest that the support value be used to assess the confidence which should be placed in individual recombinant detections of interest.

#### Accuracy of the JHMM method

The JHMM method of [4] forms a key part of our method to detect recombinants. Until now, there has not been a systematic study of the accuracy of this method. Two key outputs of this method are the locations of the inferred recombination breakpoints, and the estimated recombination parameter *ρ*. Here, we study the accuracy of these inferences for our simulated datasets.

#### Recombination breakpoints

For each recombination, we calculate the distance between the true and inferred breakpoints. For ease of comparison, we restrict this analysis to the case where each recombinant sequence has exactly two parents (one recombination), which avoids the problems of matching breakpoints in the same sequence to each other.

We find in general (see Fig 16) that the breakpoints are very accurately inferred, with 38.4% of all breakpoints inferred exactly, and 75.0% being at most 5AA from the true value. There is also a slight but noticeable positive bias, where the inferred breakpoints tend to be slightly larger than the true breakpoints (S14 Fig). This can be best explained by noting that the JHMM method infers the best (Viterbi) path from left to right, and recombinations are considered relatively unlikely; hence a recombination will tend to be inferred slightly later than it actually is, particularly if both parents’ sequences are identical around the breakpoint.

**Fig 16.**
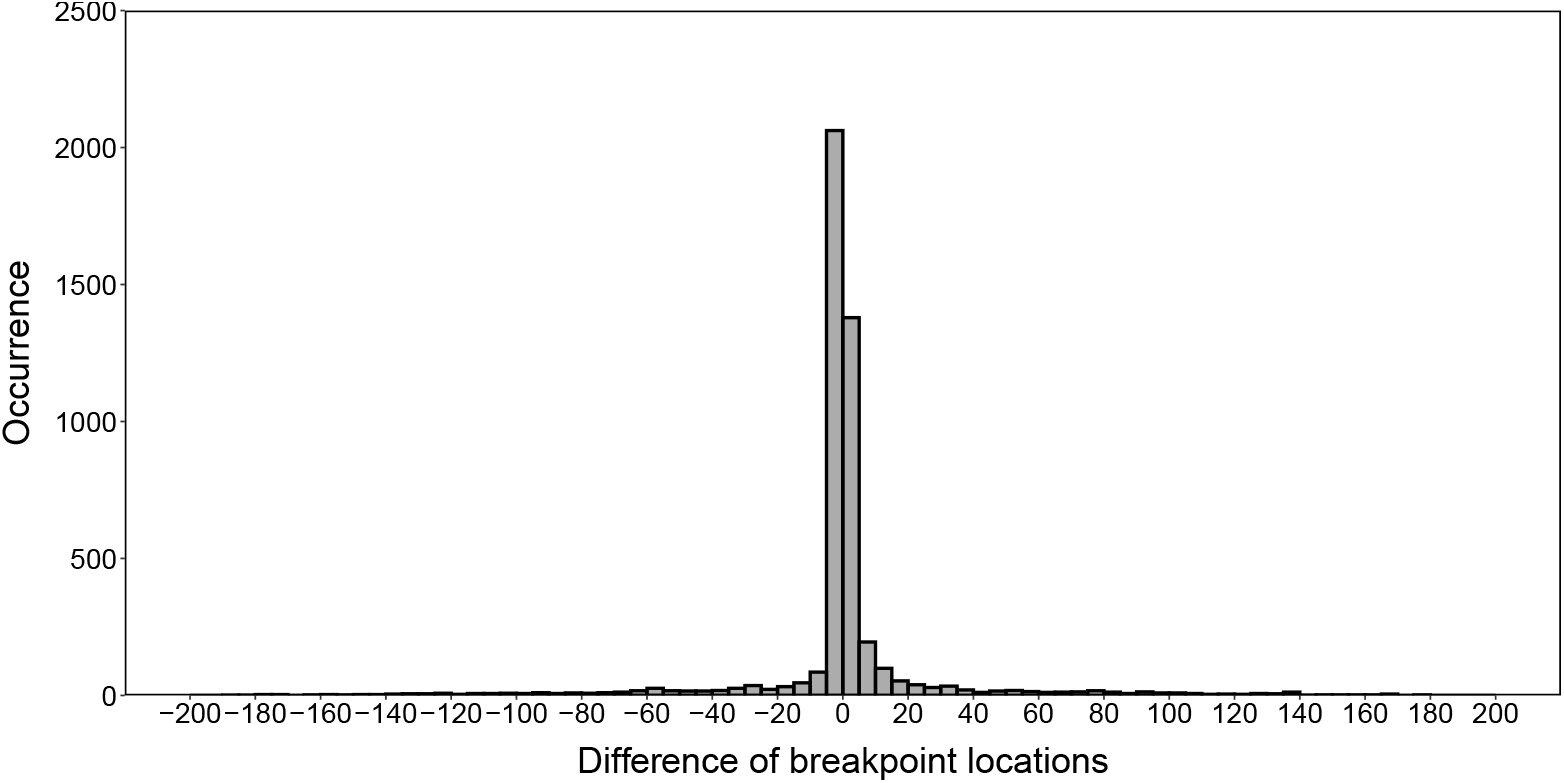
Breakpoint inference error of the JHMM method under default simulation parameters.

Finally, we note that the breakpoint accuracy appears to be very robust to indel events; this is expected, since the method explicitly accounts for these events.

#### Recombination rate

The parameter ρ is directly related to the recombination rate in the dataset (although it does not provide a rate in terms of time dimension). As such, an accurate estimate of ρ is valuable for molecular phylogeneticists. We observe in our simulated datasets (S15 Fig-S18 Fig) that the inferred values of *ρ* provide an accurate estimate of the recombinaton rate.

On the other hand, the inferred ρ can also be affected by mutation rate (Fig 17) and (to a lesser extent) indel events (S19 Fig-S20 Fig); here, an increasing rate of non-recombination events leads to some of them being mistaken for recombination, distorting the inference of the recombination rate. This indicates that the use of the JHMM to infer the true recombination rate has the potential to be inaccurate.

**Fig 17.**
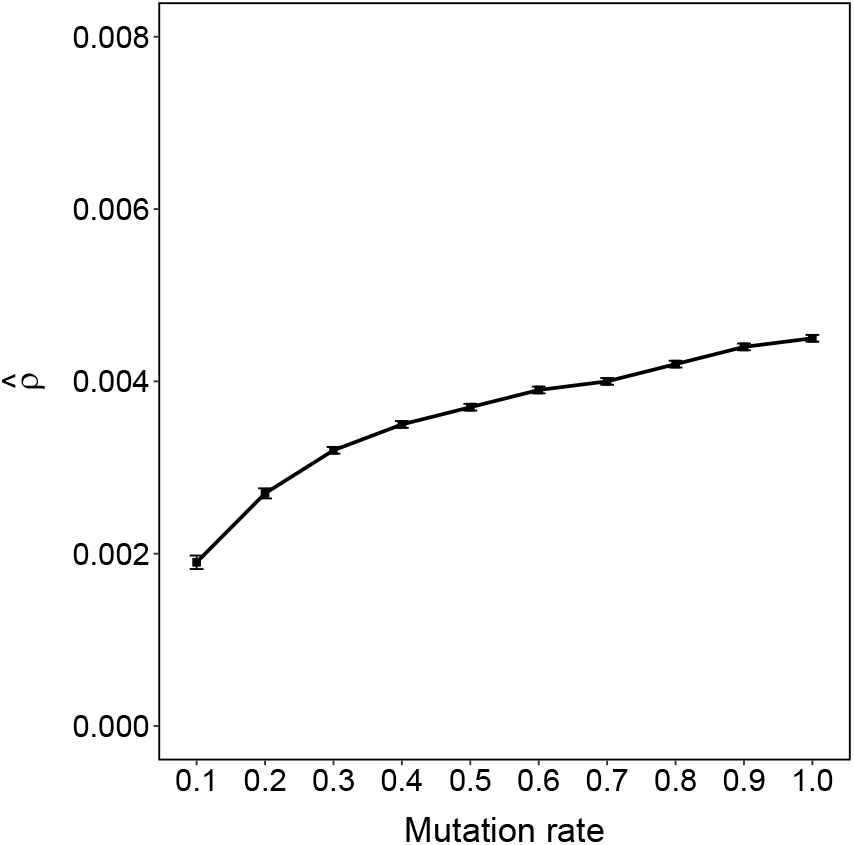
Estimated *ρ* (and 95% CI) with varying mutation rate (but constant number of recombinations).

### Discussion

In this paper, we have developed a statistical method to detect recombinant sequences from a large set of unaligned genetic sequences without a reference panel. We can also assess the reliability of the inferred recombinants with a bootstrapping-based tool. Comparisons between recombinant and non-recombinant DBL*α* types reveal a series of biologically meaningful results; for example, recombination is more frequent within ups types and DBL*α* subclasses, and non-recombinants are more conserved than recombinants. Simulations show that our method performs very well even when there is a high recombination rate, long sequences, or a large dataset. Crucially, it maintains its accuracy in the presence of insertions and deletions, where methods which require an alignment would normally fail.

We note that our method is set up to detect only recent recombinants; for example, if a more ancient recombination produces a sequence that diverges into two lineages, the lineages will be preferentially matched to each other by the JHMM, and it is possible that no recombination will be detected. Note that ‘recent’ in this context only means that the recombinant sequence has not yet diverged; it is uncertain what timescale this corresponds to. For example, although recombination events have been reported on epidemiologically relevant timescales of several years [10], a recombinant may continue to be ‘recent’ for far longer than that. The Ghana dataset studied in this paper is the first of a longitudinal dataset collected over several seasons, which may give insight into the frequency and patterns of recombination on epidemiological timescales; this is the subject of current work.

Furthermore, there is an implicit assumption that recombinations do not ‘interact’ with each other, i.e., that they are sufficiently far apart either in the evolutionary network or in the genome that we can decompose the dataset into recombinant triples and assess those independently. This is a strong (and perhaps unrealistic, in the context of genes which have a high recombination rate) assumption which we make in order to obtain a tractable algorithm. As seen from our results, we do appear to obtain good accuracy with our detections even in cases where this assumption might not hold; assessing the exact impact of this assumption on our results is also the subject of future work.

This algorithm opens up new avenues for further analysis of *var* genes. In particular, the detection of (recent) recombinants and their parents will aid in the construction of phylogenetic networks. The ability to infer such a network of *var* genes may have important implications for monitoring, intervention, and diagnosis of malaria in the future.

Finally, although our methods are motivated primarily by the highly recombinant *var* genes, our approach is not restricted to these genes, but could be used for any genes which are recombinant but lack a reliable alignment or reference panel (e.g., detecting gene fusions in the context of RNA sequencing in human cancer bioinformatics). The scalability of our method means that it will be applicable even to large datasets, thus holding great promise for broader applications.

## Supporting information

**S1 File. Recombinant proportions across isolates and catchment areas.**

**S2 File. Detection of HBs in recombinant and non-recombinant DBL**α **types.**

**S3 File. Matching recombination numbers to real data**

**S1 Fig. Distribution of source segment length in mosaic representations of Ghana data.** There is a peak of source segments which less than 5AA, which appear to be the artifacts of the JHMM method.

**S2 Fig. Proportions (and 95% confidence intervals) of recombinants for each isolate.** The horizontal dashed line displays the overall proportion of recombinant sequences in the entire dataset.

**S3 Fig. Frequency of DBL**α **types in the isolates of the Ghana dataset.**

**S4 Fig. Distribution of source segment count from the JHMM output in the Ghana data.**

**S5 Fig. The number of recombinant triples detected by our algorithm for varying sequence length.** The reference line indicates the true number of recombinant triples in the dataset.

**S6 Fig. Distribution of support values for varying proportions of recombinant sequences.** Red points represent the median of support values (same hereinafter).

**S7 Fig. Distribution of support values for varying numbers of recombinations per recombinant sequence.**

**S8 Fig. Distribution of support values for varying dataset size.**

**S9 Fig. Distribution of support values for varying sequence length.**

**S10 Fig. Distribution of support values for varying mutation rate.**

**S11 Fig. Distribution of support values for different models of amino acid evolution.**

**S12 Fig. Distribution of support values for varying indel rate.**

**S13 Fig. Distribution of support values for varying indel size.**

**S14 Fig. Breakpoint inference of the JHMM method under default simulation parameters.** Most points cluster around the line y = *x*, indicating a high accuracy of breakpoint inference. However, this is a slight positive bias in the identified breakpoint location, particularly for breakpoints which occur later in the sequence.

**S15 Fig. Estimated** p **(and 95% CI) for varying proportions of recombinant sequences.** Some CIs are too short to be visible (similarly for S16 Fig-S18 Fig. 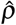 appears to grow linearly with the proportion of recombinant sequences, as expected.

**S16 Fig. Estimated** p **(and 95% CI) for varying number of recombinations per recombinant sequence.** p appears to grow linearly with the number of recombinants per sequence, as expected.

**S17 Fig. Estimated** p **(and 95% CI) for varying dataset size.** 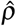 decreases slightly with increasing dataset size, although the recombination rate remains constant.

**S18 Fig. Estimated** p **(and 95% CI) for varying sequence length.** 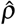 decreases in inverse proportion to the sequence length, as expected.

**S19 Fig. Estimated** p **(and 95% CI) for varying indel rate.** There is a moderate increase in *p* as indel rate increases. This is unsurprising, as some of indel events are mistaken for recombinations, distorting the inference of the recombination rate.

**S20 Fig. Estimated** p **(and 95% CI) for varying indel size.** Indel size (but constant indel rate) does not appear to have a drastic effect on estimated p.

## Data accessibility

All sequence data used in this study is available at DDBJ/ENA/GenBank: BioProject Number PRJNA396962; Accession number SAMN08902792. All the source code of proposed algorithm with test data and manuals are available from Github repository (https://github.com/qianfeng2/detREC_program).

## Acknowledgments

We would like to thank Martine Zilversmit and Mun Hua Tan for informative discussions, and David Posada for kindly providing us the source code of the recombination detection program Chimaera.

This study was supported by the Fogarty International Center at the National Institutes of Health (Program on the Ecology and Evolution of Infectious Diseases), Grant number: R01-TW009670 to KPD. Salary support for KET was provided by R01-TW009670 and The University of Melbourne. SRP was supported by a Melbourne International Engagement Award from The University of Melbourne. Salary support for MFD was provided by The University of Melbourne. QF has been supported by the China Scholarship Council. The funders had no role in study design, data collection and analysis, decision to publish, or preparation of the manuscript.

