## Supporting information for "A scalable method for identifying recombinants from unaligned sequences"

### Contents

**1 Recombinant proportions across isolates and catchment areas 3**

**2 Detection of HBs in recombinant and non-recombinant DBL$\alpha$ types 3**

**3 Matching recombination numbers to real data 4**

**4 Figures 5**

4.1 Distribution of source segment length in mosaic representations of Ghana data . 5

4.2 Proportions (and 95% confidence intervals) of recombinants for each isolate . . 6

4.3 Frequency of DBL$\alpha$ types in the isolates of the Ghana dataset . . . . . . . . . 7

4.4 Distribution of source segment count from the JHMM output in the Ghana data 8

4.5 The number of recombinant triples detected by our algorithm for varying sequence length . . . . . . . . . . . . . . . . . . . . . . . . . . . . . . . . . 9

4.6 Distribution of support values for varying proportions of recombinant sequences 10

4.7 Distribution of support values for varying numbers of recombinations per recombinant sequence . . . . . . . . . . . . . . . . . . . . . . . . . . . . . . . 11

4.8 Distribution of support values for varying dataset size . . . . . . . . . . . . . 12

4.9 Distribution of support values for varying sequence length . . . . . . . . . . 13

4.10 Distribution of support values for varying mutation rate . . . . . . . . . . . . 14

4.11 Distribution of support values for different models of amino acid evolution . . . 15

4.12 Distribution of support values for varying indel rate . . . . . . . . . . . . . . 16

4.13 Distribution of support values for varying indel size . . . . . . . . . . . . . . 17

4.14 Breakpoint inference of the JHMM method under default simulation parameters 18

4.15 Estimated $\rho$ (and 95% CI) for varying proportions of recombinant sequences . 19

4.16 Estimated $\rho$ (and 95% CI) for varying number of recombinations per recombinant sequence . . . . . . . . . . . . . . . . . . . . . . . . . . . . . . . . 20

4.17 Estimated $\rho$ (and 95% CI) for varying dataset size . . . . . . . . . . . . . . . 21

4.18 Estimated $\rho$ (and 95% CI) for varying sequence length . . . . . . . . . . . . 22

4.19 Estimated $\rho$ (and 95% CI) for varying indel rate . . . . . . . . . . . . . . . . 23

4.20 Estimated $\rho$ (and 95% CI) varying indel size . . . . . . . . . . . . . . . . . . 24

### 1 Recombinant proportions across isolates and catchment areas

We investigated the proportion of recombinants among individual isolates to determine if there were certain isolates with an elevated or reduced proportion of recombinants. Excluding isolates with less than 20 DBL$\alpha$ types [1, 2] due to limited DBL$\alpha$ type counts, resulted in a total of 158 isolates with a mean of 217.1 DBL$\alpha$ types per isolate (range 33–833). We tested if the average proportion of DBL$\alpha$ types in each isolate was equal to the overall dataset proportion with a $t$-test with a Bonferroni correction for multiple testing. There were no isolates which had a significantly different proportion of recombinants under this test (see Fig S2).

In addition, 133 isolates (82.6%) were from two catchment areas: Soe and Vea/Gowrie. A $\chi^2$ test showed no significant difference between the proportion of recombinants from these two areas ($p = 0.992$).

### 2 Detection of HBs in recombinant and non-recombinant DBL$\alpha$ types

The location of homology blocks (HBs) in each sequence was obtained using the VarDom server [3] with the default cut-off of 9.97 as a threshold to define a match. For each HB, we averaged the leftmost and rightmost relative positions of each occurrence in a sequence to obtain the overall location of the HB.

We identified in total 41 different HBs in the database (mean 5.5, range 1–10 HBs per sequence). HBs are numbered based on the frequency of occurrence [3], with HB1 the most frequent. We found that the frequency of HBs in our dataset also decreased with the numbering, with the exception of HB2 and HB3; these HBs are frequent, but lie partially outside the DBL$\alpha$ tag boundaries, making it difficult to positively identify them in the dataset. The most frequent HBs in our dataset were HB5, HB14, and HB36.

To compare sequences directly based on HBs, we used the pairwise HB similarity [4]. This is defined as the number of HBs shared between any two sequences, divided by the average number of HBs within a sequence.

#### 3 Matching recombination numbers to real data

We performed an additional simulation to match the distribution of the number of recombinations per recombinant sequence to the Ghana data. To do this, we applied the JHMM method to the Ghana data, and extracted the number of source segments matched to each target sequence (Figure S4). From this Figure, we note that it is extremely rare to have a sequence match to 8 or more source segments (i.e., 7 recombinations), so we do not allow this to happen in our simulations.

The primary difficulty here is that the JHMM method appears to slightly overestimate the number of recombinations, which means that if we simulate recombinations in exactly the same proportion as found from the Ghana data, the JHMM method produces a recombination frequency which is slightly too high. To accommodate this, we tested five sets of probabilities in simulation, and selected the probabilities of (0.02, 0.30, 0.21, 0.23, 0.14, 0.11, 0.00) for 1–7 source segments (0–6 recombinations) for each sequence. This produced a distribution of numbers of recombinations which was similar to the Ghana data.

### 4 Figures

#### 4.1 Distribution of source segment length in mosaic representations of Ghana data

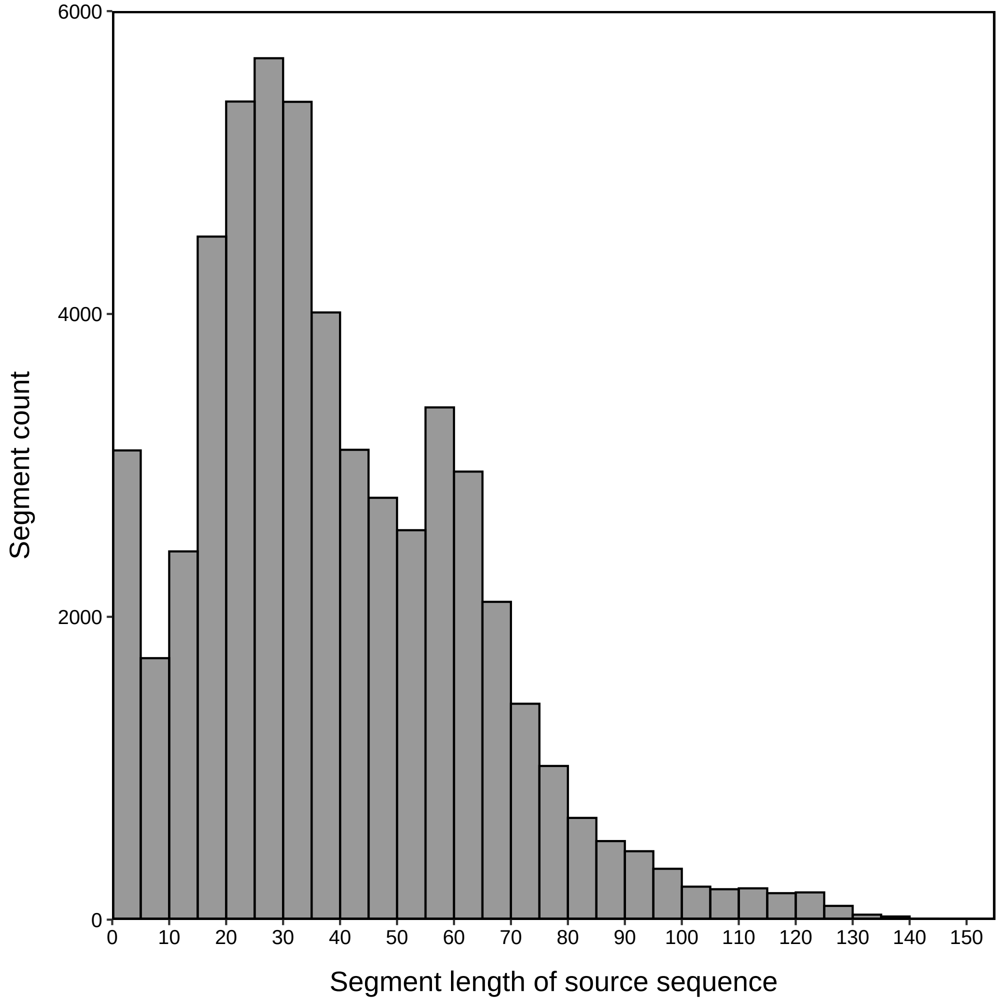


**Fig S1. Distribution of source segment length in mosaic representations of Ghana data.** There is a peak of source segments which are less than 5AA, which appear to be the artifacts of the JHMM method.

#### 4.2 Proportions (and 95% confidence intervals) of recombinants for each isolate

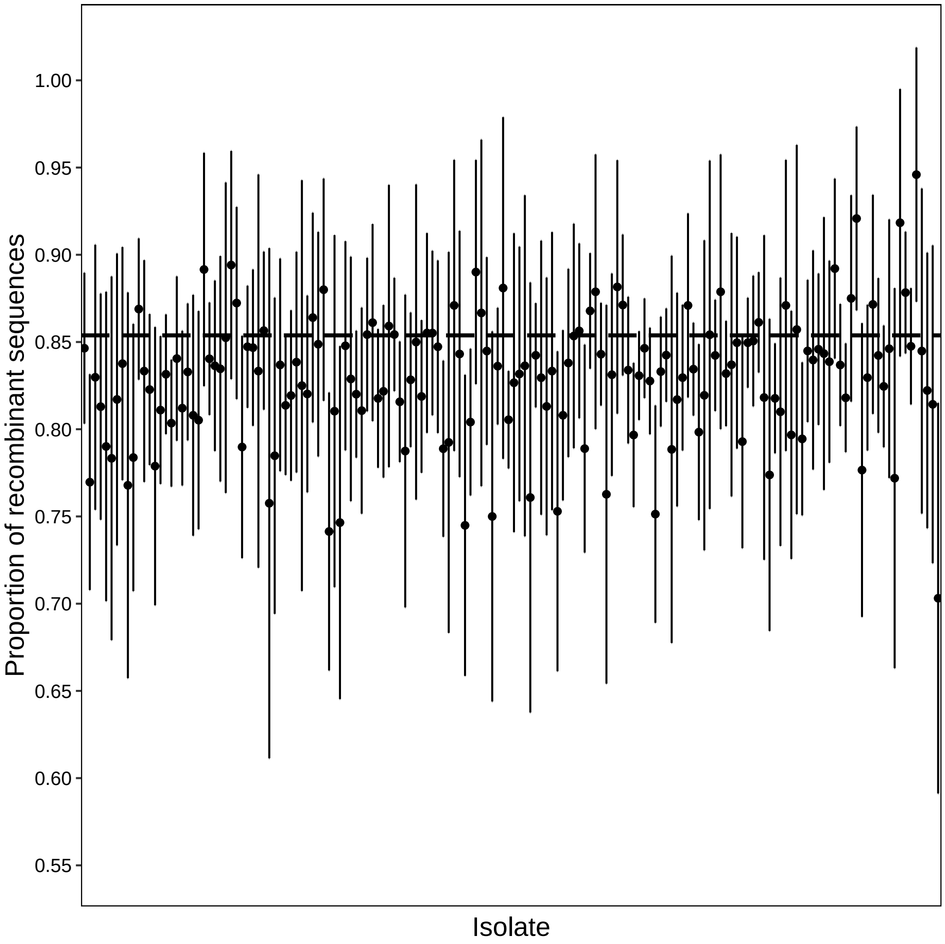


**Fig S2. Proportions (and 95% confidence intervals) of recombinants for each isolate.** The horizontal dashed line displays the overall proportion of recombinant sequences in the entire dataset.

#### 4.3 Frequency of DBL$\alpha$ types in the isolates of the Ghana dataset

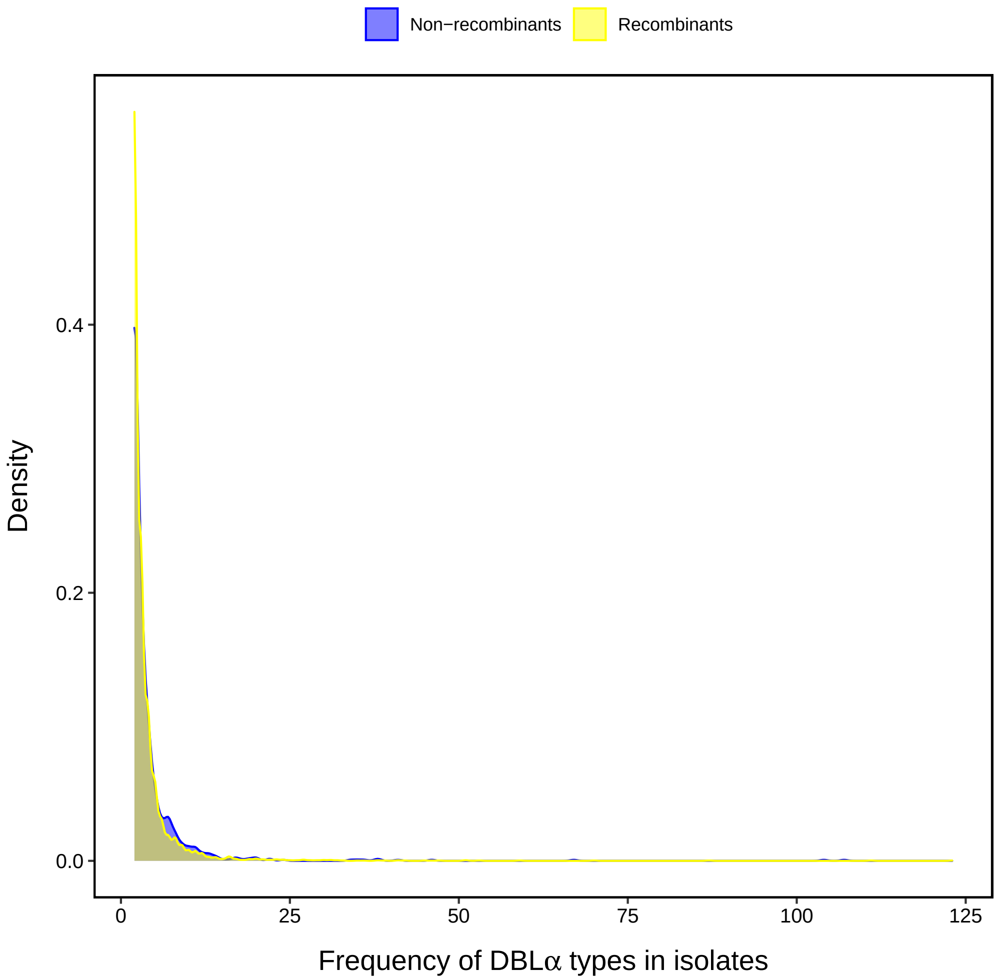


**Fig S3. Frequency of DBL$\alpha$ types in the isolates of the Ghana dataset.**

#### 4.4 Distribution of source segment count from the JHMM output in the Ghana data

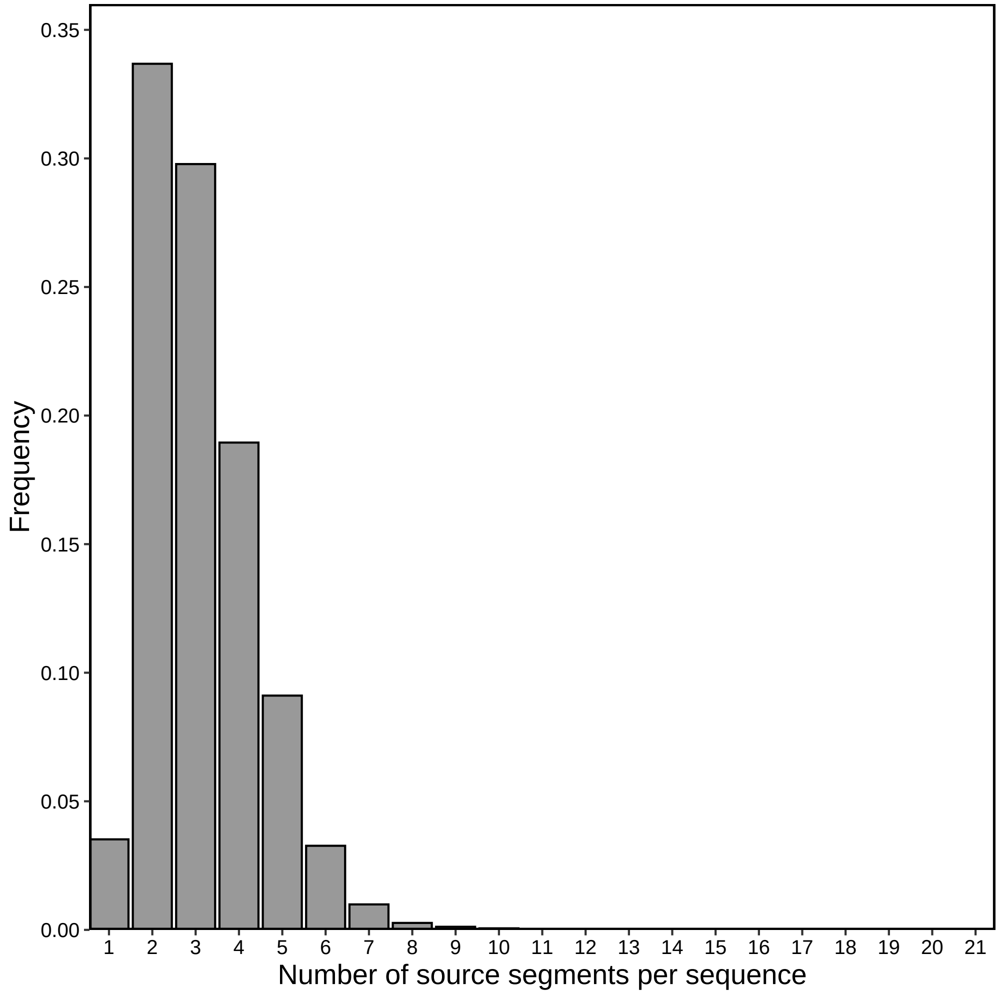


**Fig S4. Distribution of source segment count from the JHMM output in the Ghana data.**

#### 4.5 The number of recombinant triples detected by our algorithm for varying sequence length

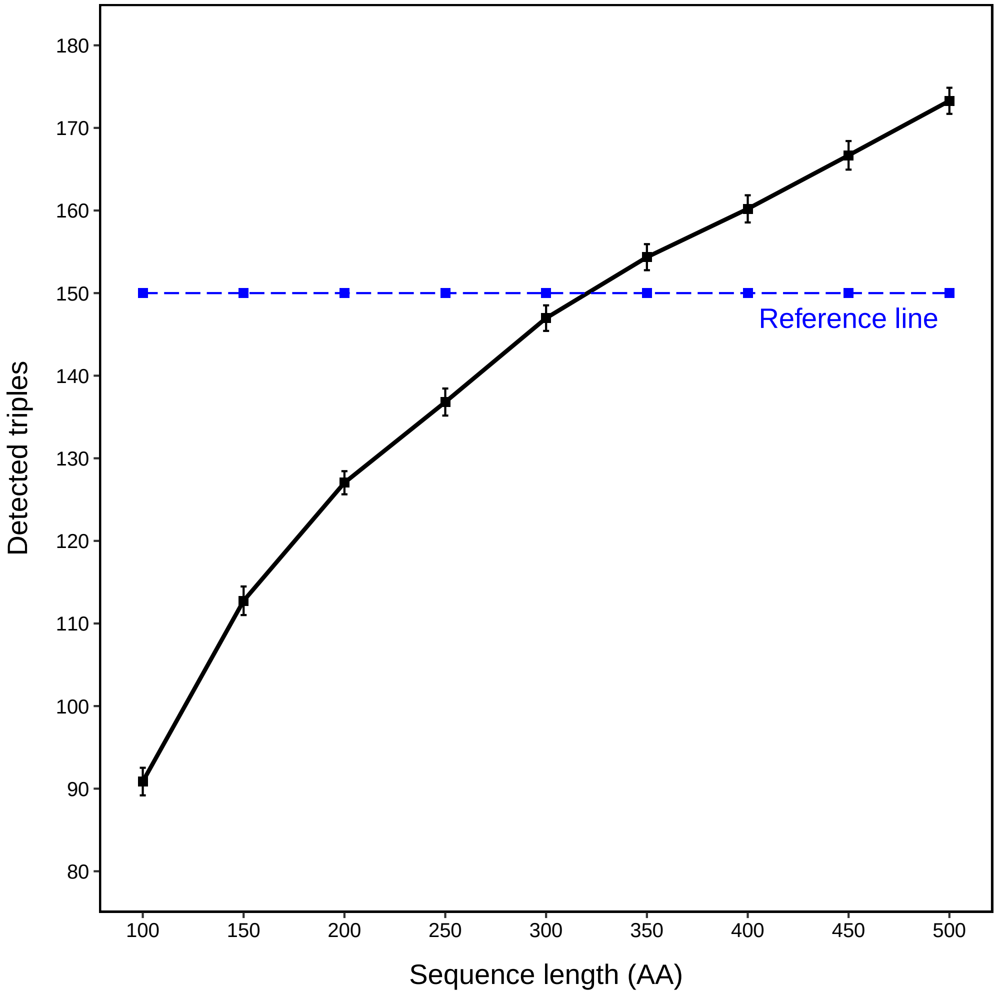


**Fig S5. The number of recombinant triples detected by our algorithm for varying sequence length.** The reference line indicates the true number of recombinant triples in the dataset.

#### 4.6 Distribution of support values for varying proportions of recombinant sequences

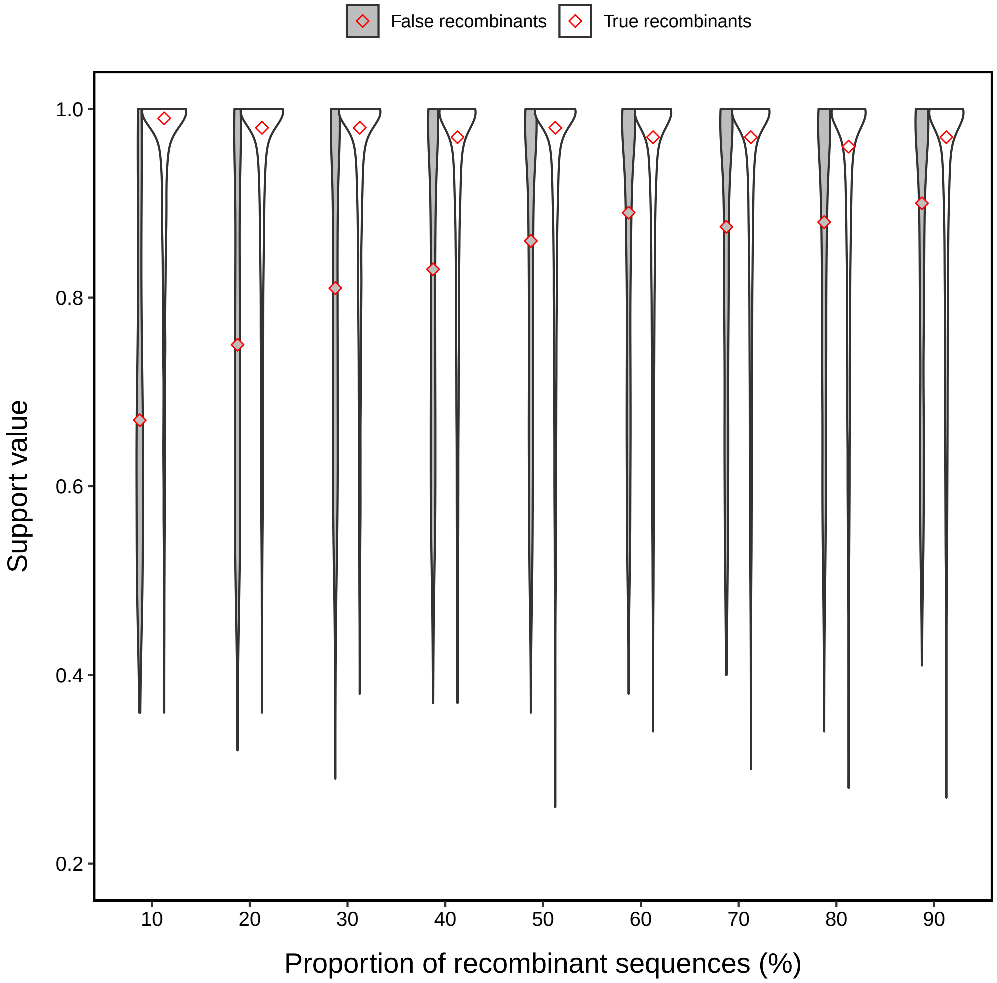


**Fig S6. Distribution of support values for varying proportions of recombinant sequences.** Red points represent the median of support values (same hereinafter).

#### 4.7 Distribution of support values for varying numbers of recombinations per recombinant sequence

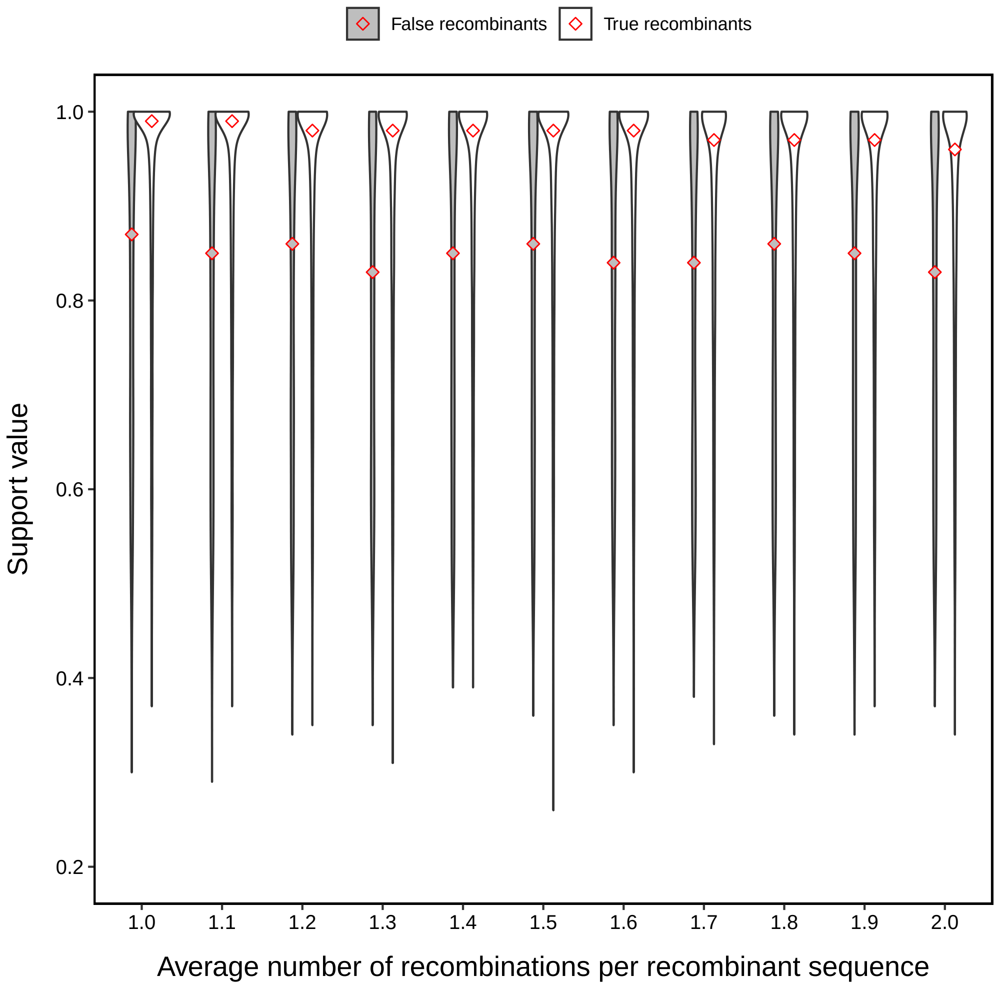


**Fig S7. Distribution of support values for varying numbers of recombinations per recombinant sequence.**

#### 4.8 Distribution of support values for varying dataset size

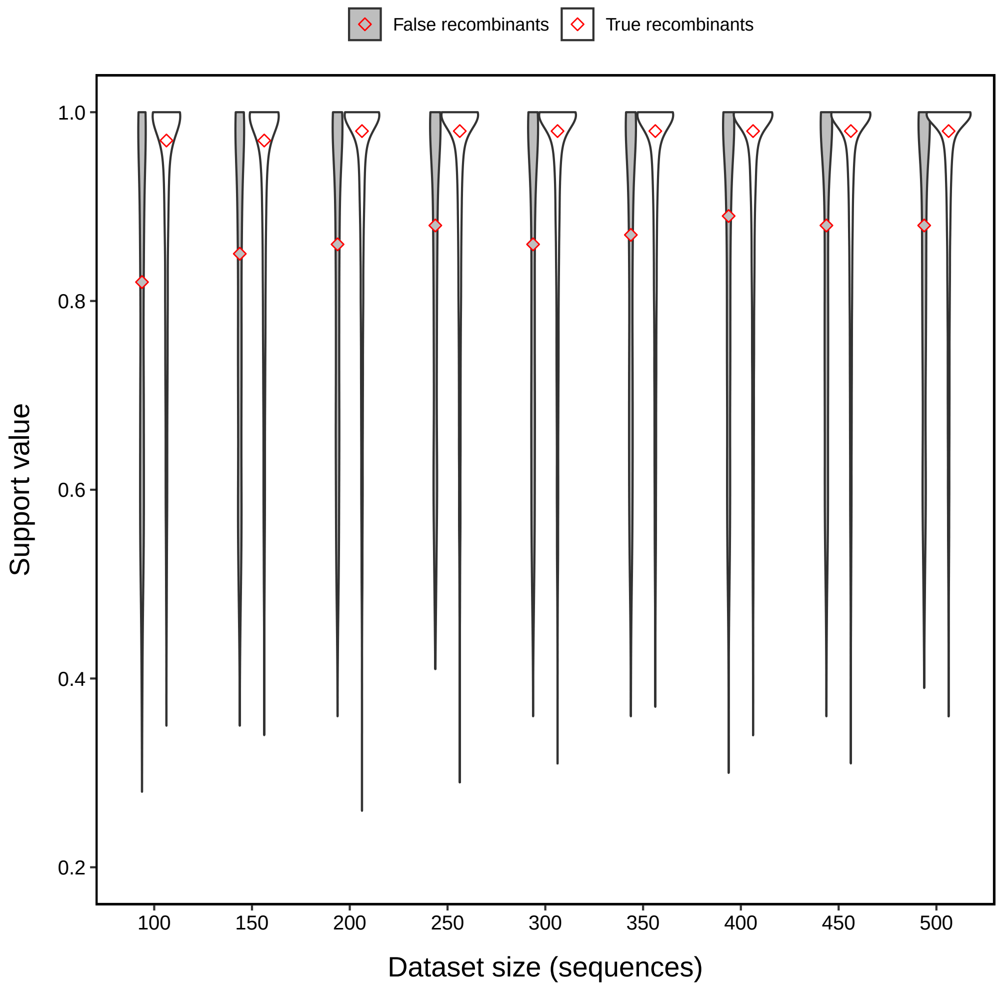


**Fig S8. Distribution of support values for varying dataset size.**

#### 4.9 Distribution of support values for varying sequence length

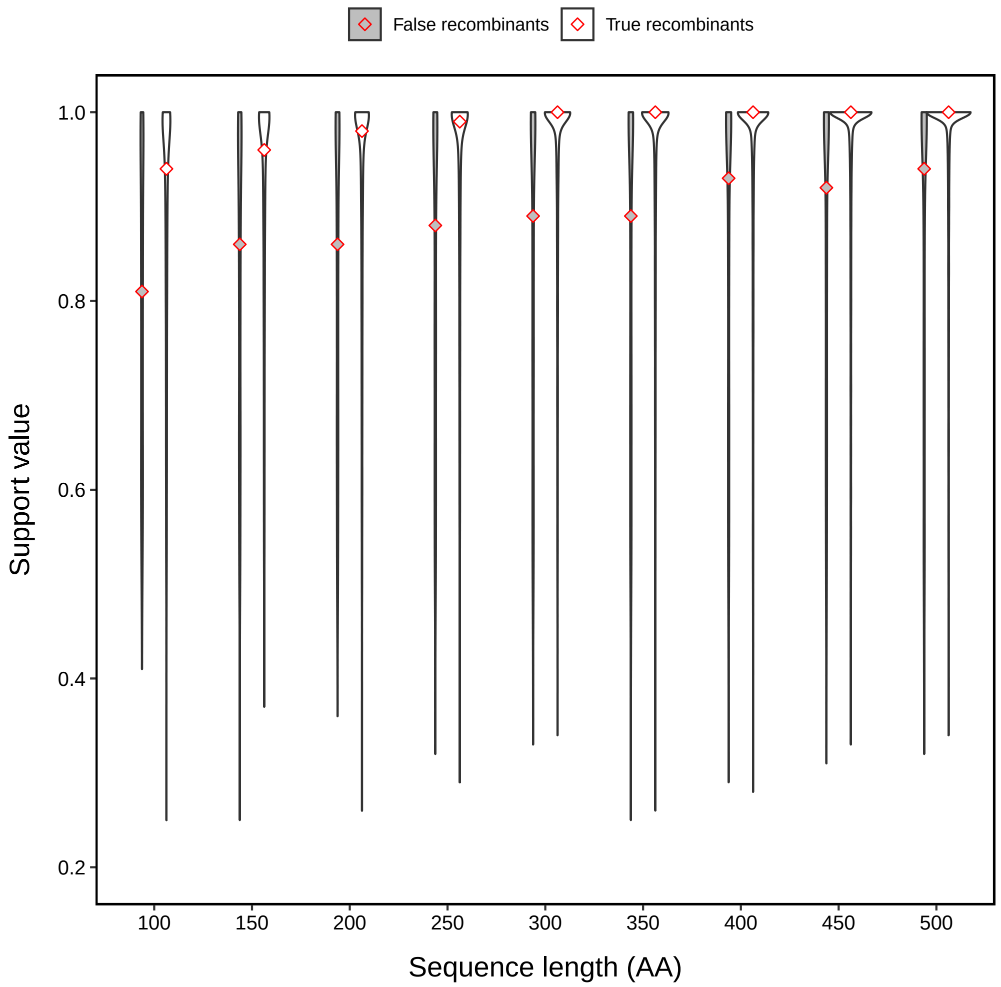


**Fig S9. Distribution of support values for varying sequence length.**

#### 4.10 Distribution of support values for varying mutation rate

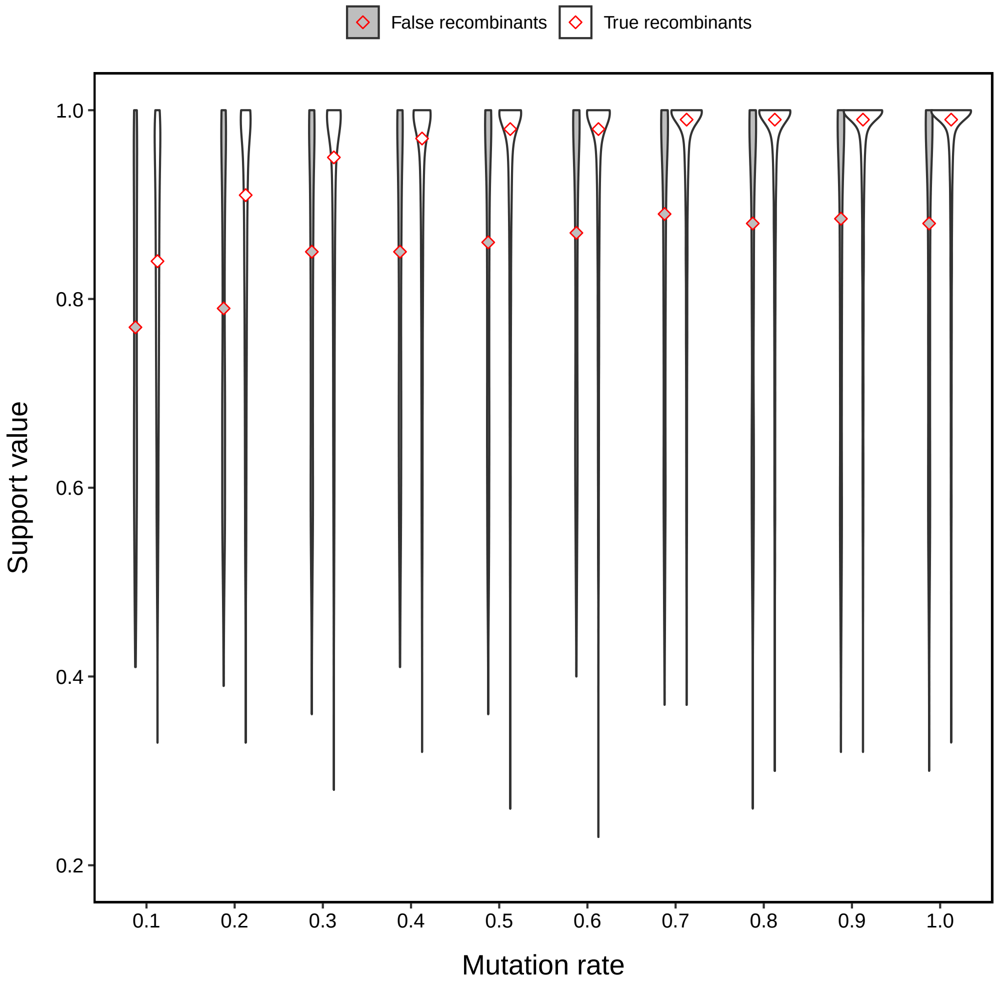


**Fig S10. Distribution of support values for varying mutation rate.**

#### 4.11 Distribution of support values for different models of amino acid evolution

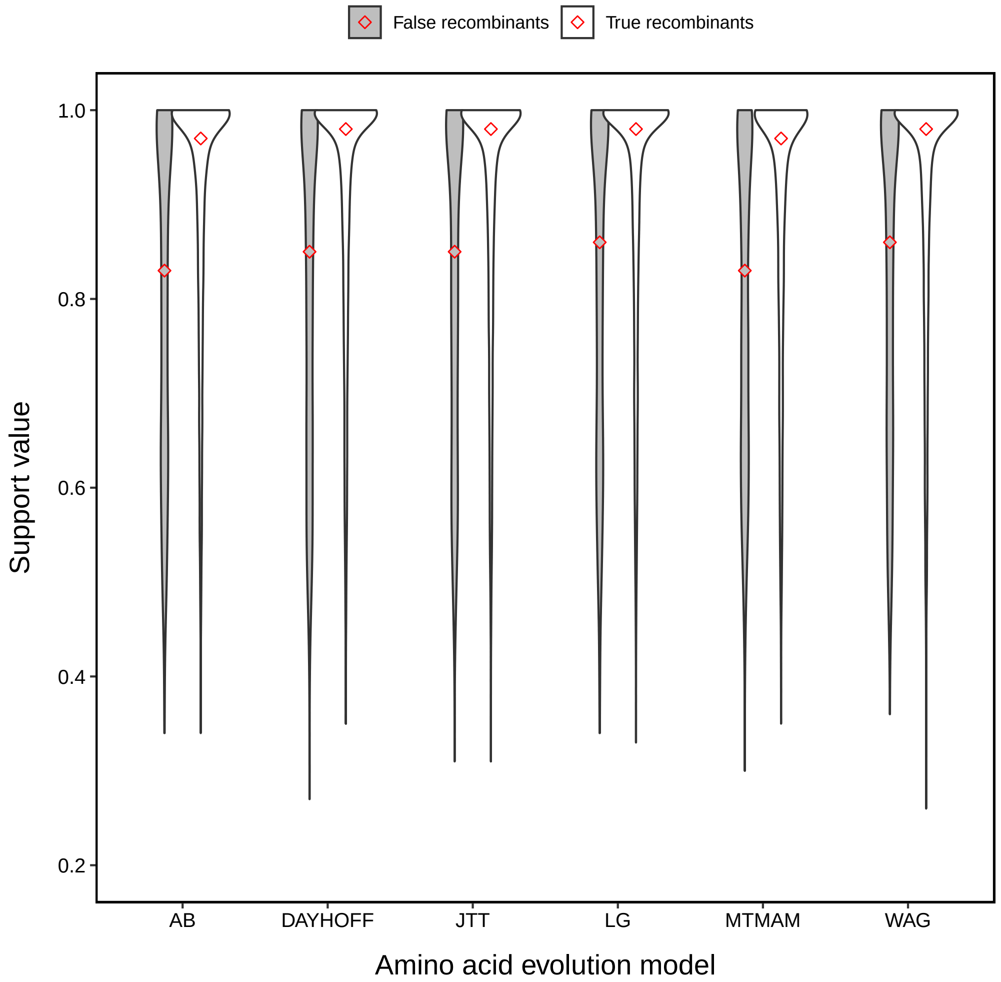


**Fig S11. Distribution of support values for different models of amino acid evolution.**

##### 4.12 Distribution of support values for varying indel rate

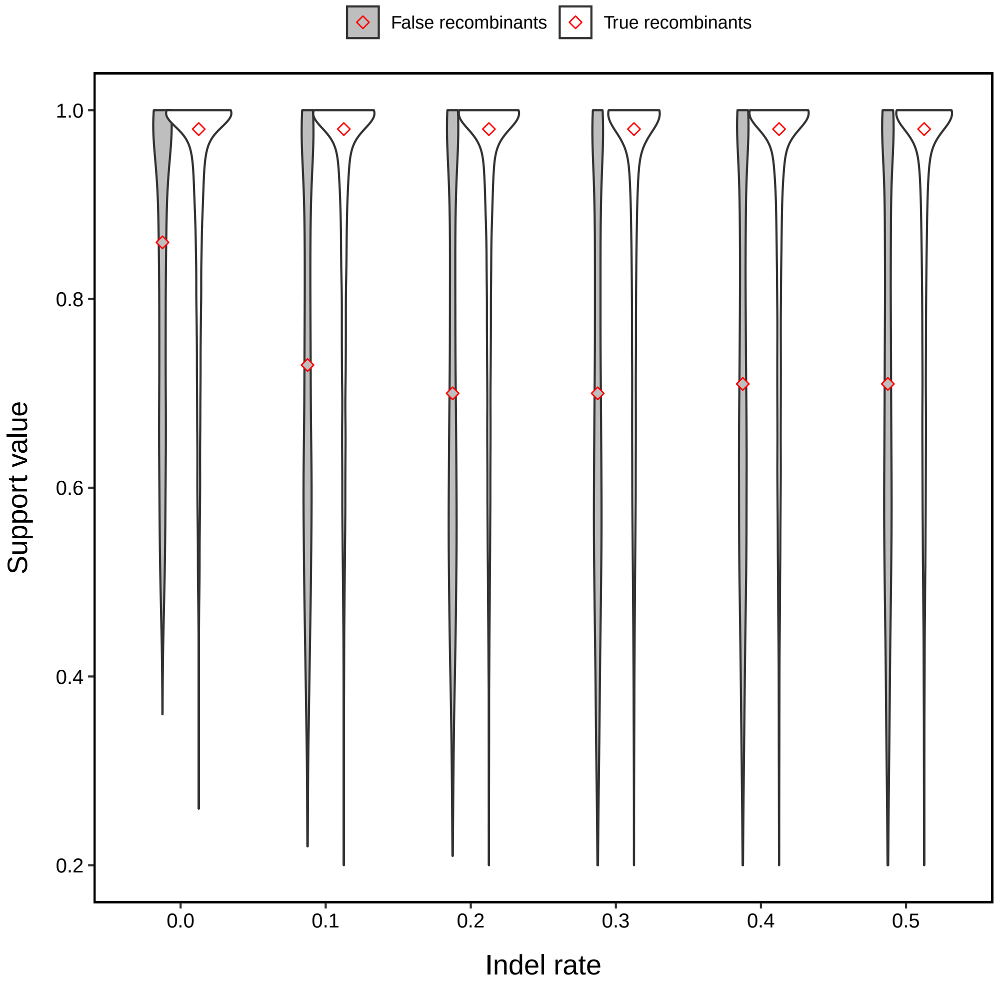


**Fig S12. Distributions of support values for varying indel rate.**

##### 4.13 Distribution of support values for varying indel size

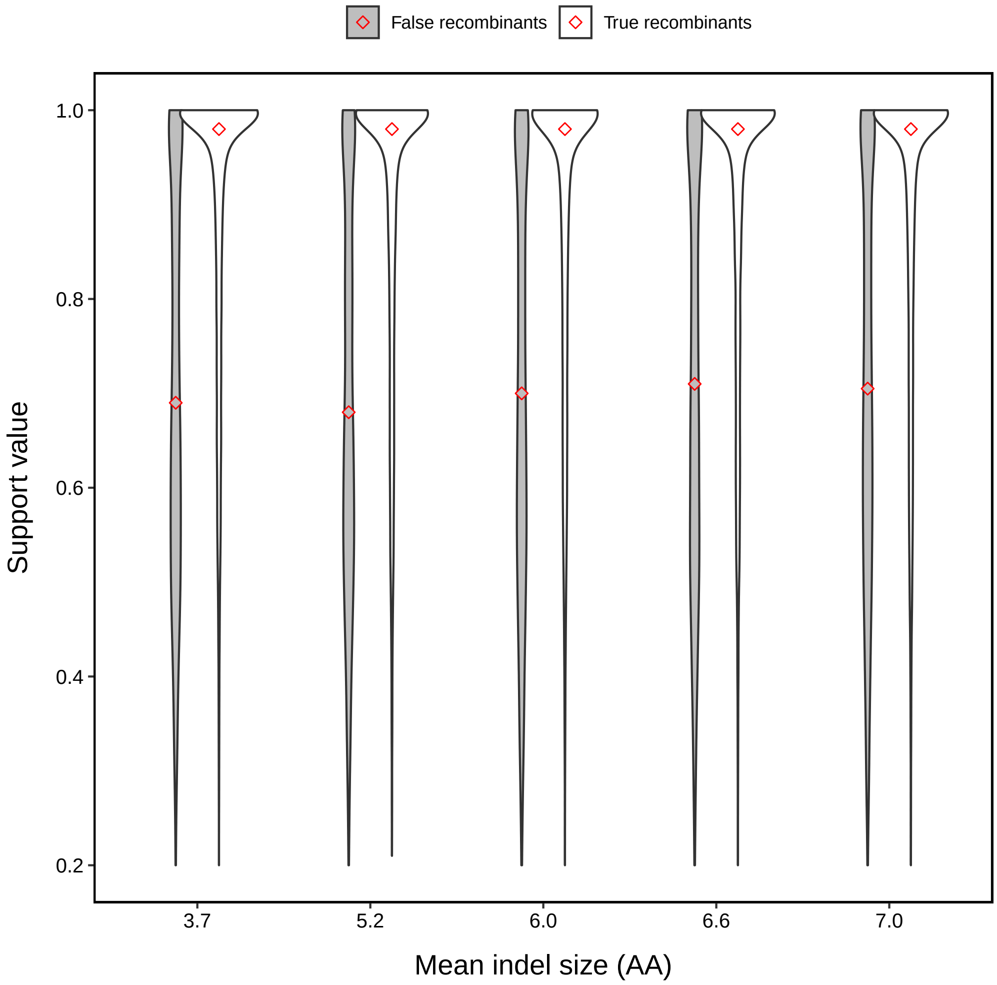


**Fig S13. Distributions of support values for varying indel size.**

#### 4.14 Breakpoint inference of the JHMM method under default simulation parameters

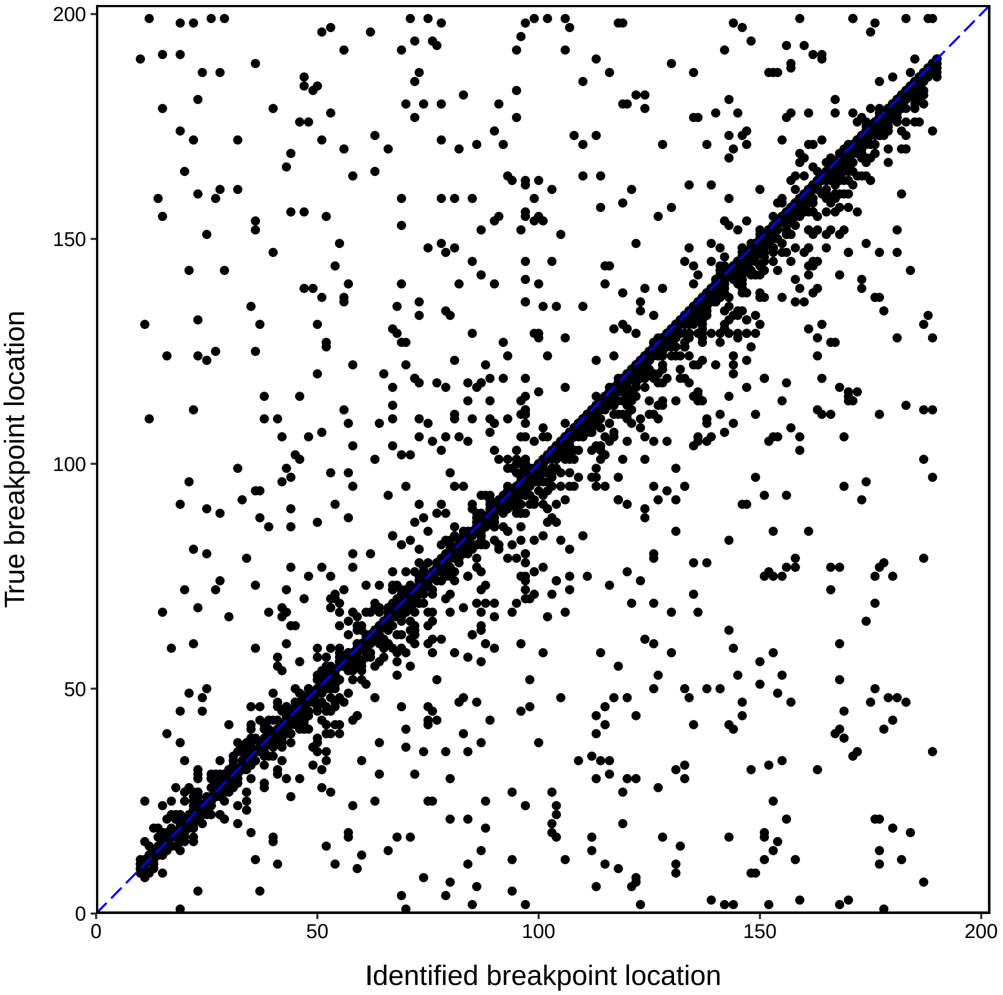


**Fig S14. Breakpoint inference of the JHMM method under default simulation parameters.** Most points cluster around the line $y = x$, indicating a high accuracy of breakpoint inference. However, this is a slight positive bias in the identified breakpoint location, particularly for breakpoints which occur later in the sequence.

##### 4.15 Estimated $\rho$ (and 95% CI) for varying proportions of recombinant sequences

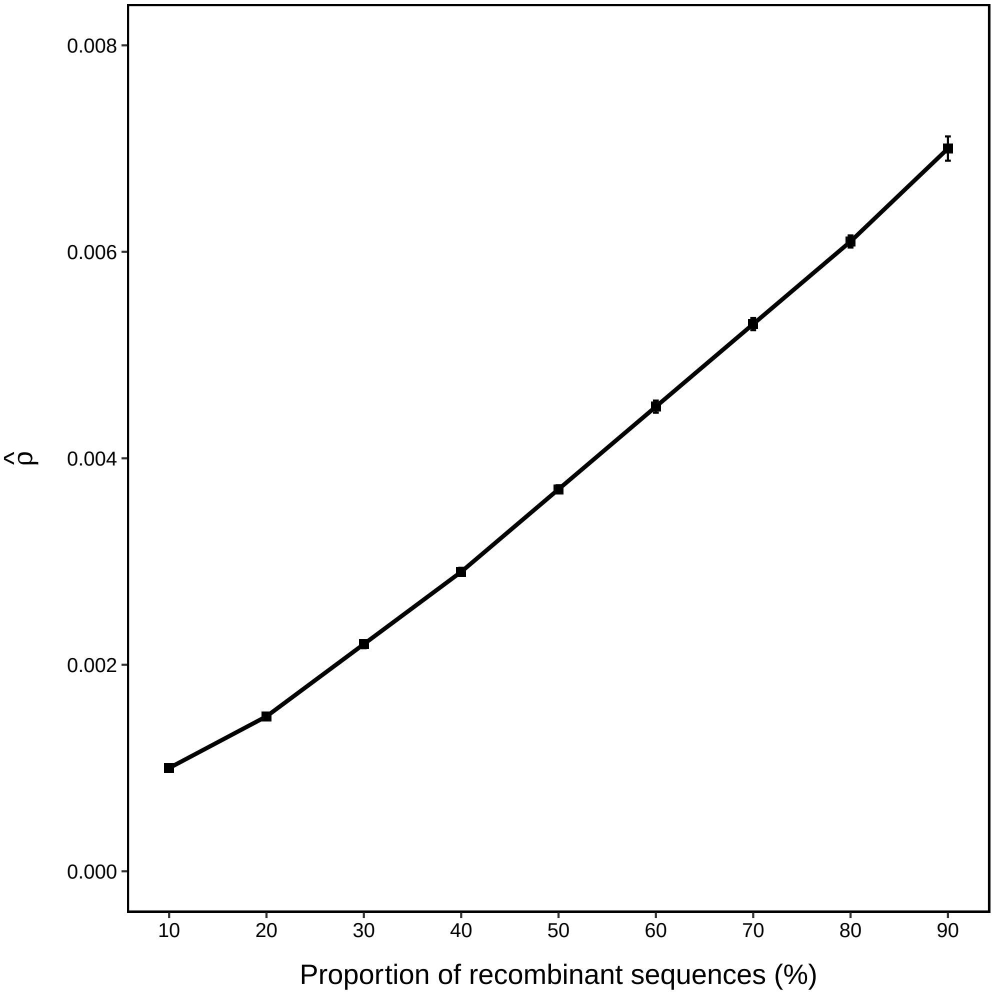


**Fig S15. Estimated $\rho$ (and 95% CI) for varying proportions of recombinant sequences.** Some CIs are too short to be visible (similarly for Figures S14–S16). $\hat{\rho}$ appears to grow linearly with the proportion of recombinant sequences, as expected.

##### 4.16 Estimated $\rho$ (and 95% CI) for varying number of recombinations per recombinant sequence

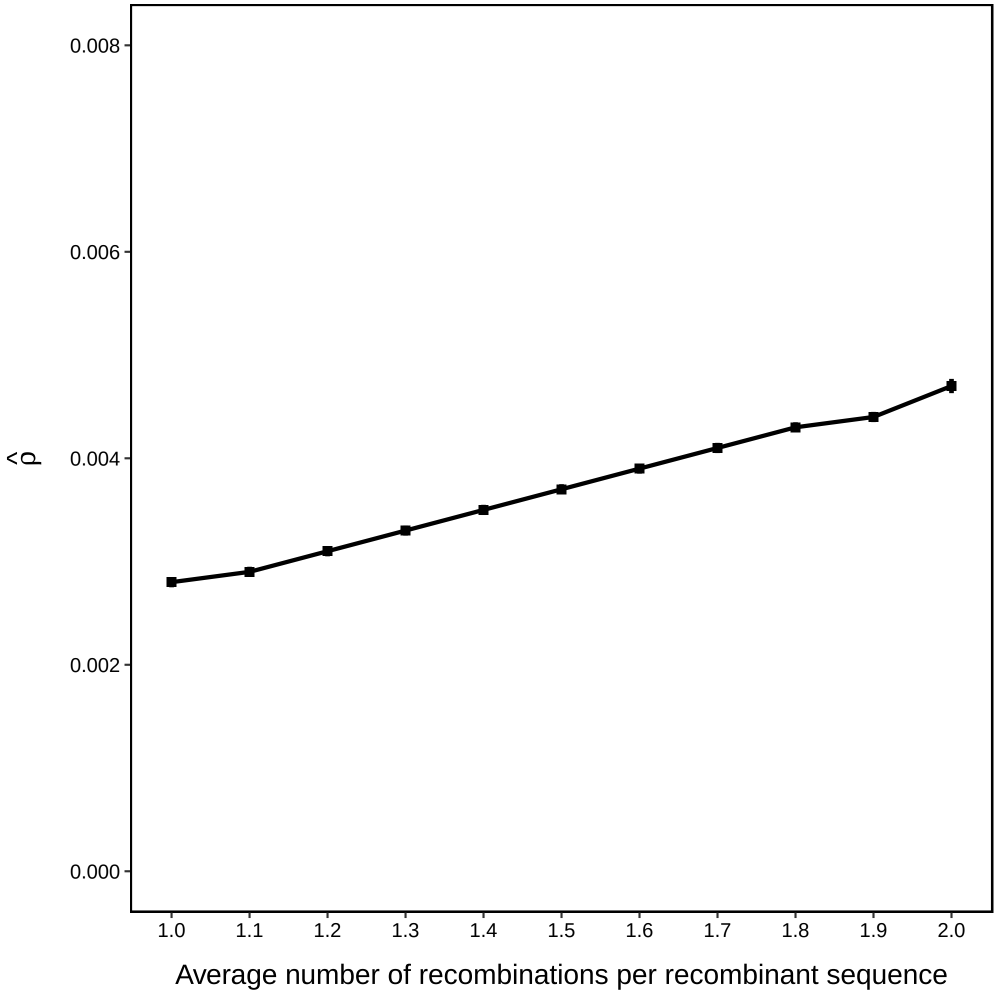


**Fig S16. Estimated $\rho$ (and 95% CI) for varying number of recombinations per recombinant sequence.** $\hat{\rho}$ appears to grow linearly with the number of recombinants per sequence, as expected.

##### 4.17 Estimated $\rho$ (and 95% CI) for varying dataset size

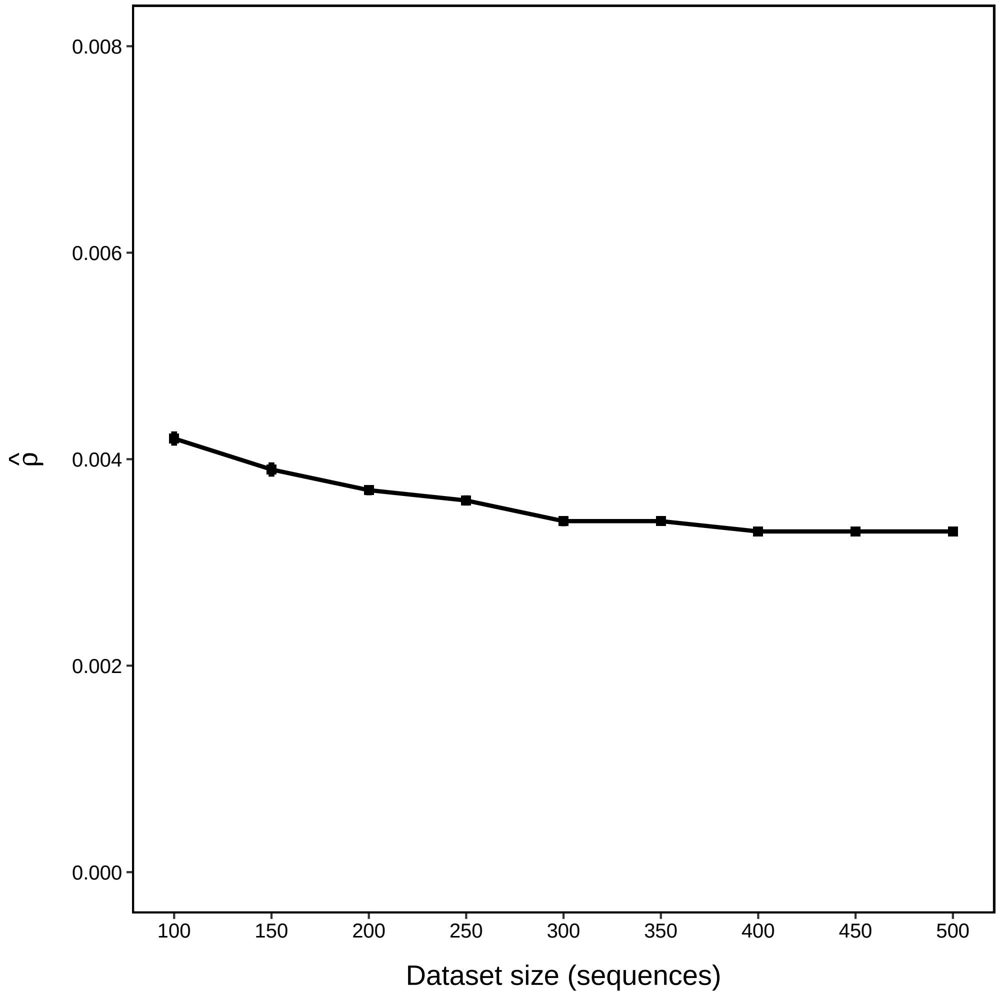


**Fig S17. Estimated $\rho$ (and 95% CI) for varying dataset size.** $\hat{\rho}$ decreases slightly with increasing dataset size, although the recombination rate remains constant.

##### 4.18 Estimated $\rho$ (and 95% CI) for varying sequence length

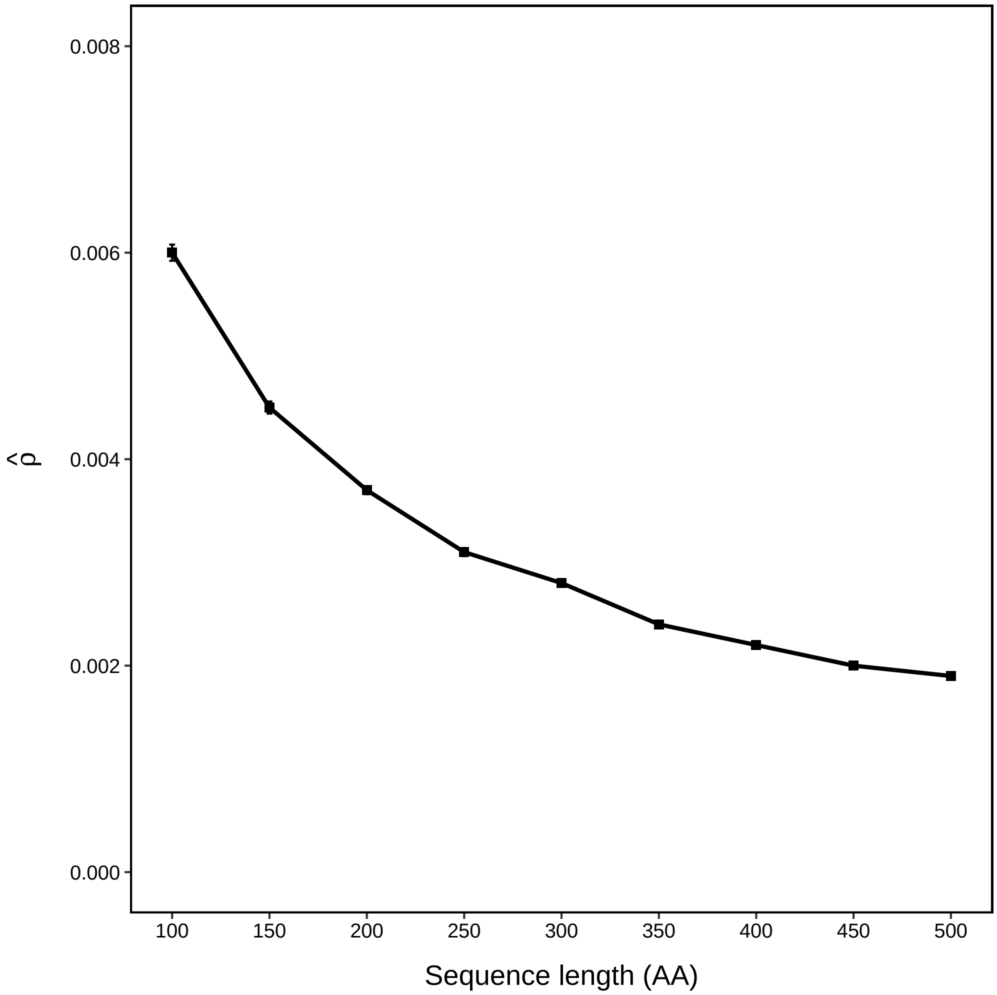


**Fig S18. Estimated $\rho$ (and 95% CI) for varying sequence length.** $\hat{\rho}$ decreases in inverse proportion to the sequence length, as expected.

#### 4.19 Estimated $\rho$ (and 95% CI) for varying indel rate

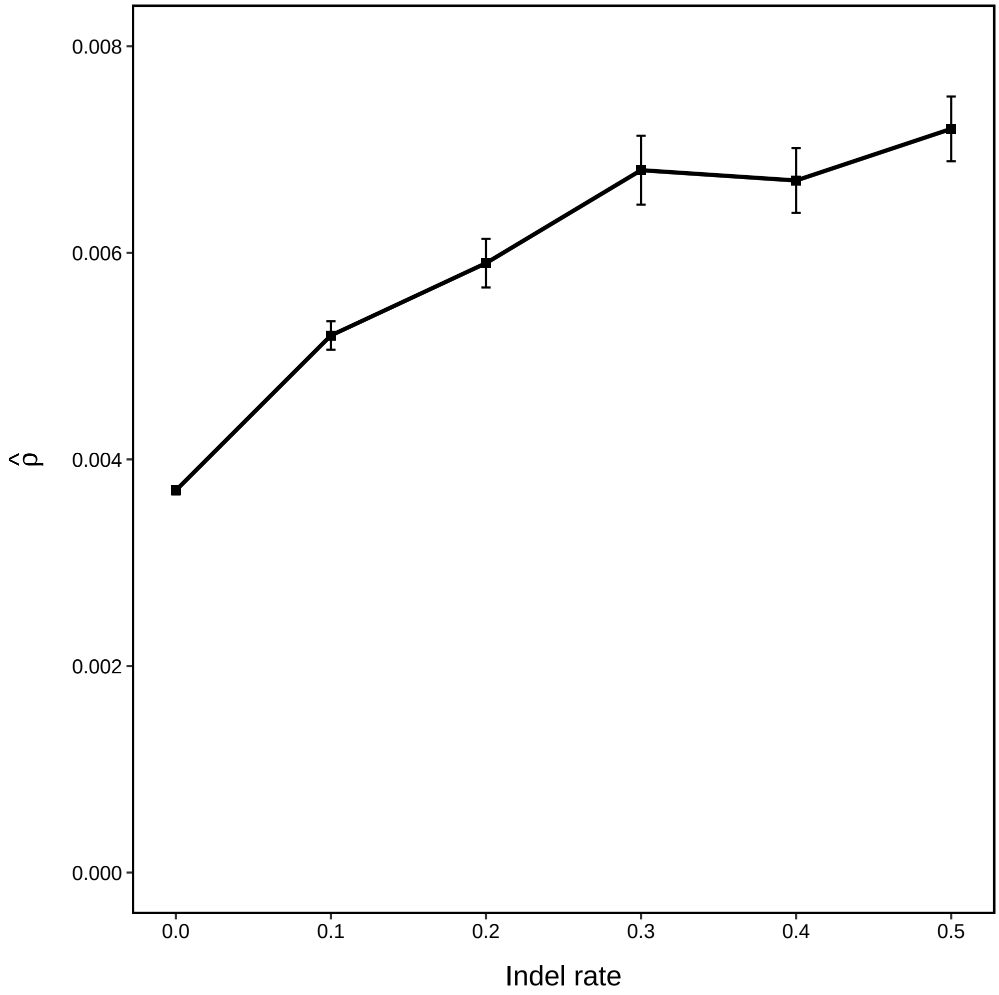


**Fig S19. Estimated $\rho$ (and 95% CI) for varying indel rate.** There is a moderate increase in $\hat{\rho}$ as indel rate increases. This is unsurprising, as some of indel events are mistaken for recombinations, distorting the inference of the recombination rate.

##### 4.20 Estimated $\rho$ (and 95% CI) varying indel size

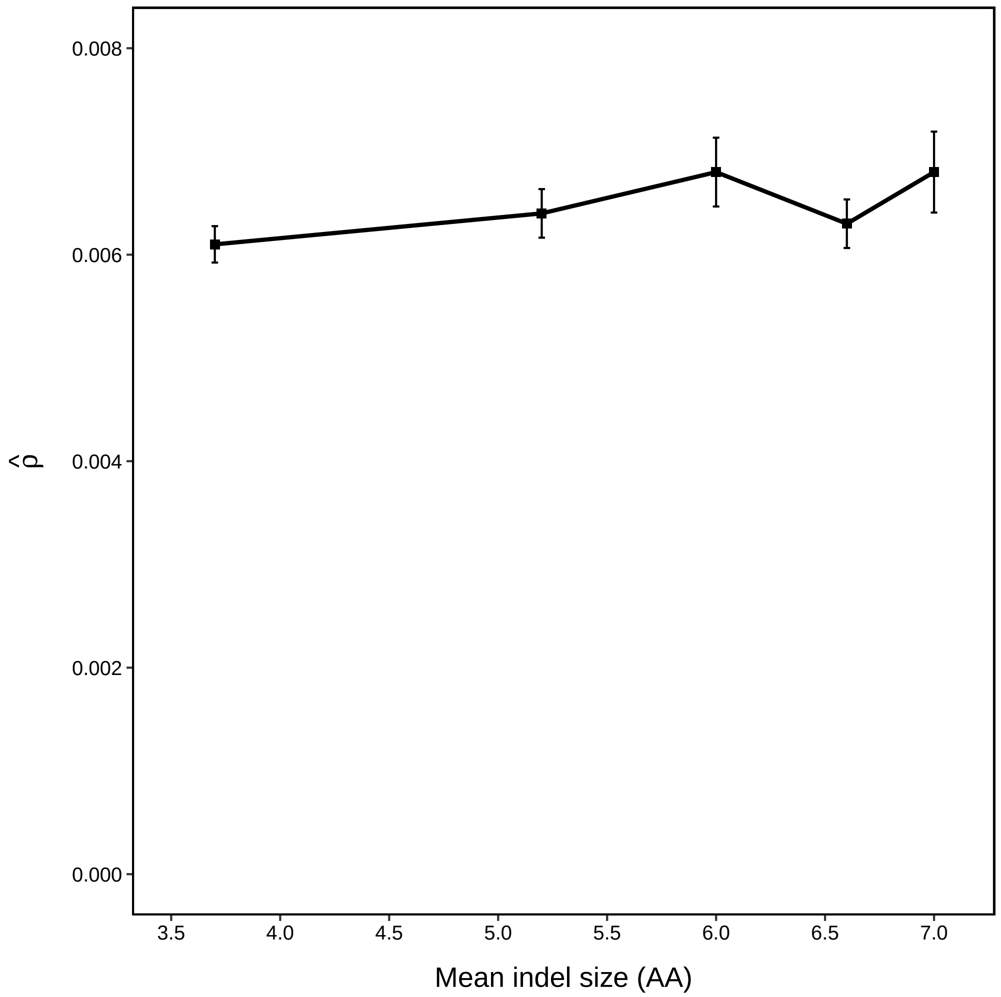


**Fig S20. Estimated $\rho$ (and 95% CI) for varying indel size.** Indel size (but constant indel rate) does not appear to have a drastic effect on the estimated $\rho$.
